# From expression footprints to causal pathways: contextualizing large signaling networks with CARNIVAL

**DOI:** 10.1101/541888

**Authors:** Anika Liu, Panuwat Trairatphisan, Enio Gjerga, Athanasios Didangelos, Jonathan Barratt, Julio Saez-Rodriguez

**Author notes:** co-first authors (Anika Liu, Panuwat Trairatphisan, Enio Gjerga).

## Abstract

While gene expression profiling is commonly used to gain an overview of cellular processes, the identification of upstream processes that drive expression changes remains a challenge. To address this issue, we introduce CARNIVAL, a causal network contextualization tool which derives network architectures from gene expression footprints. CARNIVAL (CAusal Reasoning pipeline for Network identification using Integer VALue programming) integrates different sources of prior knowledge including signed and directed protein-protein interactions, transcription factor targets, and pathway signatures. The use of prior knowledge in CARNIVAL enables capturing a broad set of upstream cellular processes and regulators, leading to a higher accuracy when benchmarked against related tools. Implementation as an integer linear programming (ILP) problem guarantees efficient computation. As a case study, we applied CARNIVAL to contextualize signaling networks from gene expression data in IgA nephropathy, a condition that can lead to chronic kidney disease. CARNIVAL identified specific signaling pathways and associated mediators dysregulated in IgAN including WNT and TGF-β, that we subsequently validated experimentally. These results demonstrated how CARNIVAL generates hypotheses on potential upstream alterations that propagate through signaling networks, providing insights into diseases.

## 1 Introduction

Cells possess a sophisticated and finely tuned signaling architecture, and its dysregulation can alter cellular behaviour leading to many diseases. A better understanding of signaling networks, therefore, allows to gain insights into disease processes and to prioritize potential targets for drug development.

Signaling networks are context specific. A network for an specific context can be inferred from dedicated data computationally. This inference can be performed based on phosphoproteomics data that directly measure key signaling players such as receptors and kinases (1, 2), preferably in combination with prior knowledge (3). However, the availability of phosphoproteomics data is often limited while gene expression data is more abundant. The inference of signaling networks based on gene expression is, therefore, an attractive approach to uncover the organisation of cellular signal transduction.

There are multiple computational tools which allow the inference of regulatory signaling networks from gene expression data. Many of these methods assume gene expression levels as a proxy for signaling protein activities and use them to construct networks (4). For instance, Huang and Fraenkel mapped transcriptomics data onto signaling pathways and then applied a Steiner’s tree algorithm for network contextualization (5). Such methods can provide valuable insight, but are limited by the fact that the abundance and activities of signaling proteins only partially correlate with gene expression (6).

To overcome this limitation, one can alternatively identify upstream signaling regulators from the profiles of downstream gene targets. One approach is to analyse gene expression footprints of signalling pathways obtained from perturbation experiments (7–9). Another one is to predict transcription factor (TF) activities based on their regulons (10, 11). However, these approaches do not provide information on the topology of signaling pathways. This information can be obtained by applying network-based approaches that can incorporate the network structure as prior information.

Given a starting prior knowledge network (PKN), upstream regulators can be inferred from downstream signaling targets in the form of a sub-network that connects direct connections and further upstream signaling events, as implemented by Melas et al. (12) (13, 14). These tools, however, only take the PKN as prior knowledge. The tool X2K, in contrast, uses expression footprint as prior knowledge to link gene expression to upstream regulatory kinases using TF and kinase enrichment, but without considering the causality of the cascades (15).

We set out to integrate the causal network approach with expression footprints to infer the whole signalling cascade. For this, we developed the causal reasoning tool CARNIVAL (CAusal Reasoning pipeline for Network identification using Integer VALue programming). CARNIVAL expands an integer linear programming (ILP) implementation for causal reasoning (12) to integrate information from TF and signaling pathway activity scoring. In addition, it can be applied not only to perturbation experiments, as in the original implementation, but also generally to compare between two or more conditions. CARNIVAL uses a comprehensive collection of pathway resources available in OmniPath as PKN (16), though other sources can be used. We performed a benchmarking study using the SBVimprover Species Translation Challenge dataset (17) and compared its performance to an alternative causal reasoning network contextualization tool CausalR (13). As a case study, we apply CARNIVAL to glomerular gene expression data on IgA nephropathy (IgAN) to gain insights on the cellular processes that regulate its pathophysiology. These were confirmed by independent experimental validation.

## 2 Methods

### 2.1 CARNIVAL pipeline

The ILP-based causal reasoning pipeline by Melas *et al.* requires a prior knowledge network, differential gene expression, as well as potential or known target(s) of perturbation for which input values are discretized. CARNIVAL incorporates several modifications: First, the objective function was customized to incorporate TF and pathway activity levels in a continuous scale. Second, dysregulated TFs are first derived with DoRothEA (7, 11) summarizing potentially noisy gene expression data into TF activities to be used as input. Last, CARNIVAL overcomes the need for known targets of perturbations which restricted the original method’s applicability. Two CARNIVAL pipelines are introduced here which will be referred henceforth as Standard CARNIVAL ‘StdCARNIVAL’ (with known perturbation targets as an input) and Inverse CARNIVAL ‘InvCARNIVAL’ (without information on targets of perturbation), see Figure 1.

**Figure 1:**
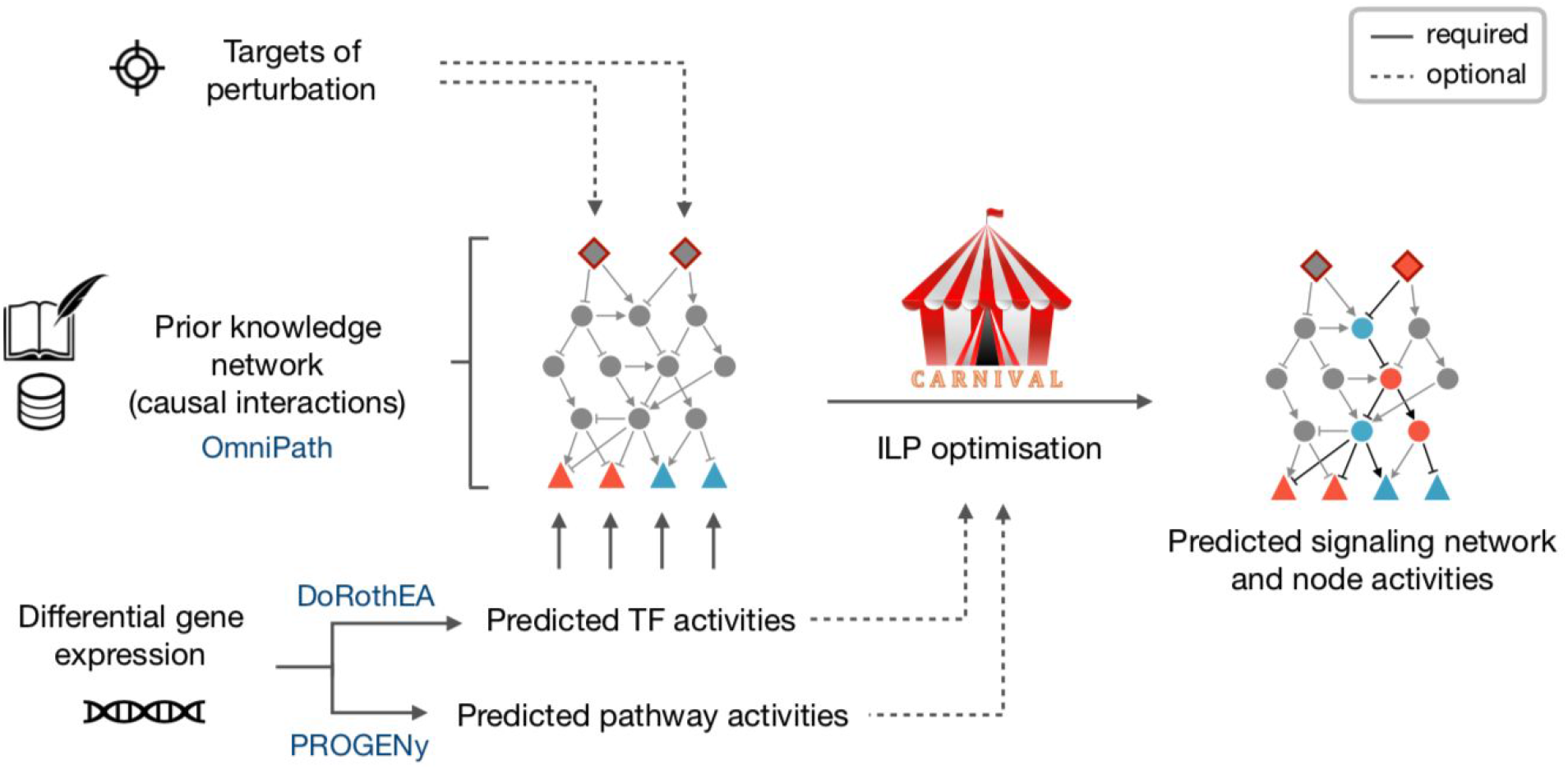
CARNIVAL pipeline. The CARNIVAL pipeline requires as input a prior knowledge network and differential gene expression. The information on perturbed targets and their effects can be assigned (StdCARNIVAL) or omitted (InvCARNIVAL). The differential gene expression is used to infer transcription factor (TF) activities with DoRothEA, which are subsequently discretized in order to formulate ILP constraints. As a result, CARNIVAL derives a family of highest scoring networks which best explain the inferred TF activities. Continuous pathway and TF activities can be additionally considered in the objective function.

### 2.2 CARNIVAL inputs

As a prior knowledge network, we applied a signed and direct human signaling network retrieve from Omnipath. The network contains 9,306 signed and directed edges connecting 3,610 nodes pooled and curated from multiple resources including Signor, Reactome and Wikipathways (16).

For TF activity prediction, we applied DoRothEA version 2 (11) which provides a framework to estimate TF activity from the gene expression of its direct target genes. The provided regulon set was filtered to include only the 289 TF-regulons with at least ten TF-target gene interactions with medium to high confidence (confidence score A, B and C as defined in (11)). Subsequently, the differential gene expression t-values processed by the *limma* R-package (18) and the filtered DoRothEA regulon were passed to the viper function in the *VIPER* package (10) to perform an analytic Rank-based Enrichment Analysis (aREA). The activities of each transcription factor in the form of normalized enrichment score (NES) were then derived from the rank of the genes and the top 50 TF scores were used as the input in CARNIVAL.

To calculate predicted pathway activity, the 14 PROGENy pathway signatures were obtained from (7, 19) and applied to differential expression t-values from *limma* (18). Based on an empirical null distribution generated through 10,000 times gene-wise permutation and the percentile corresponding to the observed value, the significance score (termed ‘score’ henceforth, Equation 1) was derived.

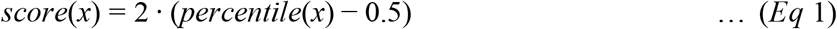

### 2.3 ILP implementation, objective function and parameter settings

We implemented the causal reasoning ILP formulation of Melas *et al.* (Suppl. Text S1) in R. The objective function is defined by Equation 2:

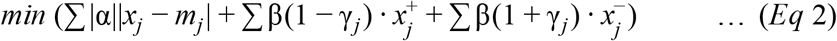

where the parameter α refers to the mismatch penalty, β to the node penalty, and the newly introduced γ to the node penalty adjustment (Equation 2).

In the previous work of Melas *et al.*, the objective function prioritizes the network in which the node activities (*x*_*j*_) explain the corresponding observed discretized measurements (*m*_*j*_) while the overall number of nodes in the network is minimized through the sum of activities 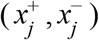 for each node *j* in the network. In CARNIVAL, we introduce the effect of inferred TF and pathway activities to adjust this tradeoff and model selection (Equation 2). For TF scores, we applied a TF-specific mismatch penalty α corresponding to the NES derived from DoRothEA. The node parameter β can then be manually assigned to scale the importance of node penalty relative to TF-scores. For pathway scores, a minimal set of representative downstream nodes was chosen for each PROGENy pathway to capture all known signal transduction routes involved while avoiding overlapping information between pathways, and with TF predictions (Suppl. Table S1). The node penalty is sign-adjusted through the γ weights which corresponds to PROGENy significance scores (Equation 1) ranging from −1 to 1. This means that the anticipated direction corresponding to pathway score is penalized less in the expected direction while more in the counterpart.

Regarding parameter settings, we implemented in this study several options to retrieve alternative top scoring solutions through the available CPLEX parameters. The solutions within 0.01% tolerance with regard to the best solution were accepted (mip pool relgap = 0.0001) and the most aggressive search strategy was employed (mip pool intensity = 4). We generated up to 500 solutions (mip limits populate = 500) fulfilling the pre-defined criteria in the solution pool and took the 100 most diverse solutions generated within one hour for further analysis (mip pool capacity = 100; mip pool replacement = 2; time limit = 3600 seconds). We applied the default setting for all other CPLEX parameters.

Additional information regarding parameter settings and the ILP problem formulation for InvCARNIVAL can be found in Suppl. Text S2. Summarized results from the study of multiple α-to-β ratios to assess parameters’ robustness can be found in Suppl. Text S3.

### 2.4 Benchmark dataset

The benchmark dataset was taken from the SBVimprover project which contains perturbations on normal human bronchial epithelial cells (17). Gene expression was measured at 6 hours after perturbation (E-MTAB-2091) in a processed form (log2 expression after GC robust multiarray averaging). Probe IDs were mapped to HGNC gene symbols and multiple entries were summarized by the mean value. Batch effects were removed using the combat function of the *sva* R-package (20). Differential gene expression was then computed with the *limma* R package (18).

The measurements with 19 phosphoprotein-binding antibodies were mapped to 14 differential protein activities using the curated regulatory sites in the PhosphositePlus knowledgebase (21). Given that only a small fraction of the PKN nodes is reported as dysregulated in CARNIVAL, the overlap between dysregulated nodes and measured protein activities was low and not suited for statistical testing.

### 2.5 Kidney datasets

Microarray data on glomerular gene expression in IgAN patients and healthy living donors (HLD) were obtained from 5 publicly accessible studies (22–25), see details in Suppl. Table S2. Study- and platform-dependent batch effects were mitigated using the combat function from the *sva* R-package (20), and differential gene expression is determined with the limma R-package (18).

### 2.6 CausalR

The CausalR package identifies dysregulated nodes and networks by scanning for nodes with sign-consistent shortest paths to the observations (13). With the SCAN (Sequential Causal Analysis of Networks) method, path lengths from one to five edges are scanned and potentially dysregulated nodes are identified which constantly score among the top 150 based on the number of explained observations. Matched observations increase the score (+1), mismatched ones decrease it (−1), and unmatched or ambiguously matched nodes are not included in the scoring.

### 2.7 Two-step inference approach to KEGG pathways attribution

In our study, we assume that the inferred node activities from CARNIVAL represent upstream signaling and should hence map well with the KEGG pathways attributed to the corresponding perturbation. Hence, a two-step inference approach was developed for validation (Figure 2). In the first step, we assume that an over-representation of up-regulated CARNIVAL nodes indicates higher activity pathway and, conversely, down-representation represents down-regulation. Up- and down-regulated pathways were predicted with a hypergeometric test from the *Category* package in R on the dysregulated nodes inferred by CARNIVAL. The universe in this regard was set to all nodes present in the PKN derived from Omnipath and the curated KEGG pathway sets were obtained from MSigDB (26). A significance test was only performed if at least one set member of the pathway was present in the given CARNIVAL node set.

**Figure 2.**
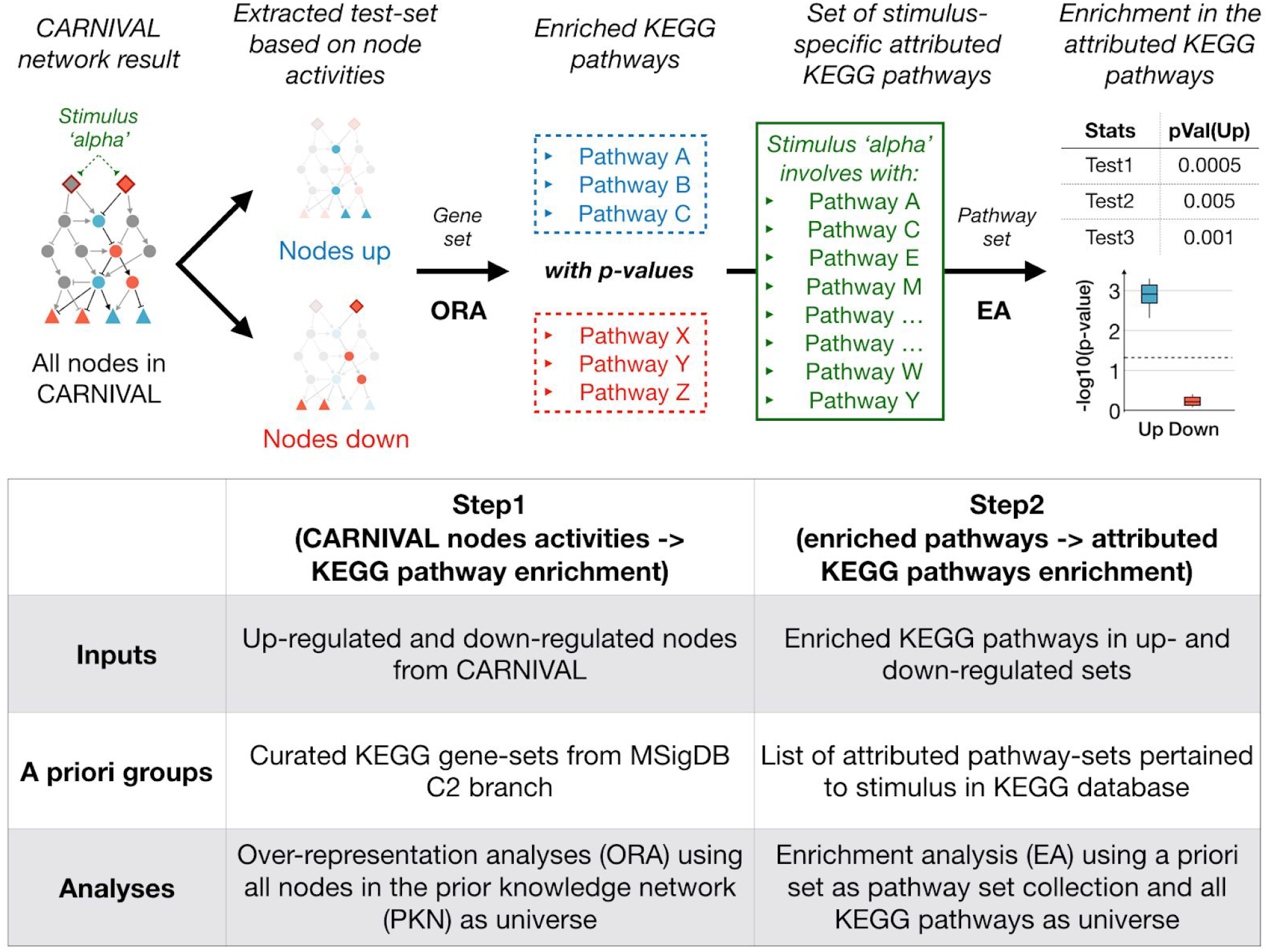
Two-step inference analysis to determine whether relevant molecular processes were identified in CARNIVAL. First, dysregulated pathways were inferred by over-representation of the nodes in CARNIVAL solution networks based on the KEGG pathway sets in MSigDB. In the second step, an enrichment analysis was performed on the identified dysregulated pathways using stimulus-specific pathways as prior set. The distributions of p-values from multiple statistical tests are reported as final result. A significantly enrichment of the attributed pathways in the direction that the target protein is perturbed is expected.

In the second step, we assumed that the list of KEGG pathways attributed to the perturbation should be up-regulated upon perturbation while others are not. We then check if the pathways identified with CARNIVAL fulfill this assumption (Figure 2). Specifically, we evaluate if both the up- and down-regulated pathways with their FDR-adjusted p-values from the first step were enriched in the attributed pathways. Here, we applied the function *runGSA* with the *stouffer*, *tailStrength*, *wilcoxon and reporter* methods in the *piano* R-package (27). The mean and standard deviation across the four methods of the resulting p-values were reported. A Gene Set Enrichment Analysis (GSEA) was applied to gene expression directly to identify a baseline performance.

### 2.8 Network topological analyses and text mining

To identify central nodes with regulatory features, we computed quantitative measures from topological analyses of CARNIVAL networks and compare them to the ones derived from randomized networks. These measures were extracted using the functions *degree, betweenness, hub_score and authority_score* in the *igraph* R-package (28). The average nodes’ activities over the range of size penalty parameter (β in (0.03; 0.1; 0.3; 0.5; 0.8)) in the CARNIVAL networks were applied as weights for the calculation of these topological analyses measures. The R-package *BiRewire* (29) was applied to generate 100 randomized networks with preserved distributions of node degrees as well as interaction signs. Student’s t-test was subsequently used to determine significant differences between the distributions of network topology measures from CARNIVAL networks versus the ones from randomized networks. To assess the “novelty” of genes identified in this analysis in the context of IgA nephropathy we queried PubMed Central using the approved gene symbol and approved gene name from the HUGO gene nomenclature committee database (genenames.org), together with the following keywords: “IgA nephropathy”, “IgAN”, “IgA glomerulonephritis” and “Berger’s disease”. All search terms were enclosed in quotations to ensure that terms were searched in full and retrieved abstracts were examined to exclude erroneous matches.

## 3 Results

### 3.1 Benchmarking on the SBV improver dataset

To evaluate the performance of the CARNIVAL pipeline, we applied it to the SBVimprover Species Translation Challenge dataset which provides phosphorylation and gene expression data for multiple perturbations (17). We applied both the StdCARNIVAL and InvCARNIVAL pipelines to evaluate the effect of information from the perturbation targets onto the resulting networks. The results from both pipelines were then compared to the ones generated by CausalR as well as GSEA.

This study provides phosphoprotein data, that in principle lends itself to validate the estimated activity of nodes with CARNIVAL. However, only 3-to-4 out of 19 phosphosites in the phosphoprotein dataset could be mapped to CARNIVAL node activities per condition, which is insufficient for statistical analyses (see Methods). We therefore used an alternative validation. We first determined which pathways are known to be linked to the perturbations by different molecules according to the KEGG database (30); referred to perturbation-attributed pathways henceforth. We then performed an enrichment analysis to define whether they are more up- or down-regulated with a two-step inference approach (see Methods and Figure 2). This gives insights into how well expected pathways and their regulatory direction are captured by CARNIVAL.

#### 3.1.1 Incorporation of predicted TF and pathway activities

The normalized enrichment scores (NES) from DoRothEA were used as an estimate for the degree of dysregulation (see Methods). For StdCARNIVAL, significant enrichment of the perturbation-attributed pathway set in up-regulated pathways is only achieved for IL1-β (IL1B) and TGF-α (TGFA) with the introduction of TF weights (Suppl. Figure S1). In InvCARNIVAL, where the targets of perturbations are not known, the results with pathway weights showed a significant enrichment of perturbation-attributed pathways in up-regulated pathways for PDGF-β (PDGFB), IL1-β, EGF, TGF-α, and flagellin while only TGF-α and FSL1 was significant without pathway weights (Suppl. Figure S2). TGF-α was more enriched in activated than in inhibited pathways with pathway weights but this trend was inverse for IGF2 and AREG. While being imperfect, PROGENy weight still provides an overall improvement in detecting more dysregulated pathways in the up-regulated direction (5 versus 2). Given that improved performance was found with the implementation of TF and pathway weights, these were implemented in our subsequent benchmarking.

#### 3.1.2 Comparison of StdCARNIVAL, InvCARNIVAL, GSEA and CausalR

To benchmark CARNIVAL results against related tools, we applied the two-step inference approach also on their results and made comparisons. As an overview, the perturbations of the significant up-regulated sets found in InvCARNIVAL (PDGF-β, IL1-β, EGF, TGF-α, and flagellin) were also identified in StdCARNIVAL (Figure 3; Suppl. Figure S3). In contrast, NTF3 and FSL1 only showed a significant enrichment of the up-regulated gene sets in StdCARNIVAL. This suggests that a wider coverage of pathways can be detected by StdCARNIVAL, where the perturbation target is known. The same number of enrichment of activated pathways in the perturbation-attributed pathways (up-regulated) captured with InvCARNIVAL is slightly higher than the one with pathway inference from differential gene expression directly (n=5 versus n=4). TGF-α and IL1-β were captured with both approaches, while IFN-γ (IFNG) and TNF-α (TNFA) were only significantly enriched in activated pathways inferred by GSEA, and IL1-β, PDGF-β and EGF by InvCARNIVAL. However, the trend of correct directionality, i.e. significant in the up-regulated gene sets and insignificant in the down-regulated gene sets, was only captured with InvCARNIVAL. CausalR captured a significant enrichment of activated pathways in the perturbation-attributed pathway set for IFN-γ, IL1-β and TNF-α, showing an equal performance to InvCARNIVAL in terms of detecting correct directionality but still captured less numbers of up-regulated pathways than GSEA and Std/InvCARNIVAL. (Figure 3).

**Figure 3.**
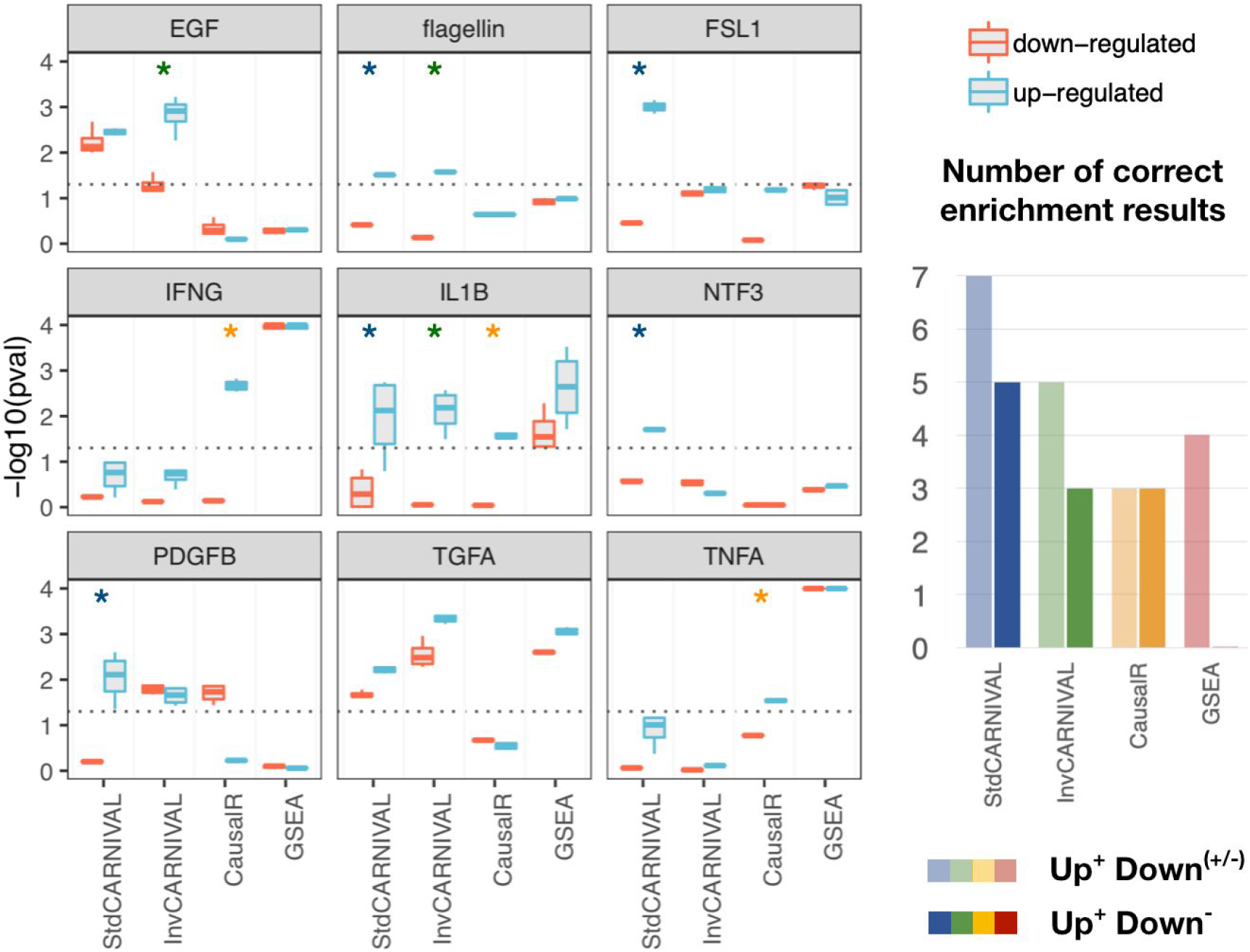
Comparison of the enrichment results of the perturbation-attributed pathway set in dysregulated pathways inferred with different tools. An enrichment of the perturbation-attributed pathway set among the significant pathways was determined. The significance level of 0.05 is indicated by the dotted lines. Asterisks (*) indicate the expected correct results where the enrichment results are significant in the up-regulated set and insignificant in the down-regulated set.

We then compared the network topology of the CARNIVAL versus CausalR networks. We found a bias towards hub nodes for CausalR but not for CARNIVAL. The degree distribution of edges in CARNIVAL was very similar to the one of the prior knowledge network (PKN), see Suppl. Figure S4.

### 3.2 Inferring signaling networks in IgAN

Immunoglobulin A nephropathy (IgAN) is a common chronic kidney disease (CKD) accounting for 35% of all renal transplantations in adults (31) and is the most frequent form of glomerulonephritis. It is characterized by the deposition of aberrant IgA in the kidney’s glomeruli, structures of blood vessels involved in blood filtration. The pathogenic IgA-containing immune complexes trigger the activation of inflammation and fibrosis (32). Further improvement in early diagnosis and treatment of IgAN are needed and can only be achieved by a better understanding of the disease’s mechanisms.

#### 3.2.1 InvCARNIVAL results

In this study, we generated the causal networks from the differential gene expression in glomeruli between groups of healthy subject versus IgAN patients. Given that the node penalty did not affect the performance but might result in minor fluctuations, this analysis was performed with different node penalties to achieve more robust results (β in (0.03; 0.1; 0.3; 0.5; 0.8), see Methods and Figure 4). Given that the adherens junction set is the most dysregulated one, the set members are highlighted in the network. Thereby, only one transcription factor (TCF7) inferred by DoRothEA is represented in this gene set, while 13 associated nodes and four input nodes are solely inferred by causal reasoning with CARNIVAL. Hence, CARNIVAL was able to capture more pathway members and their connections than through TF-regulons alone.

**Figure 4.**
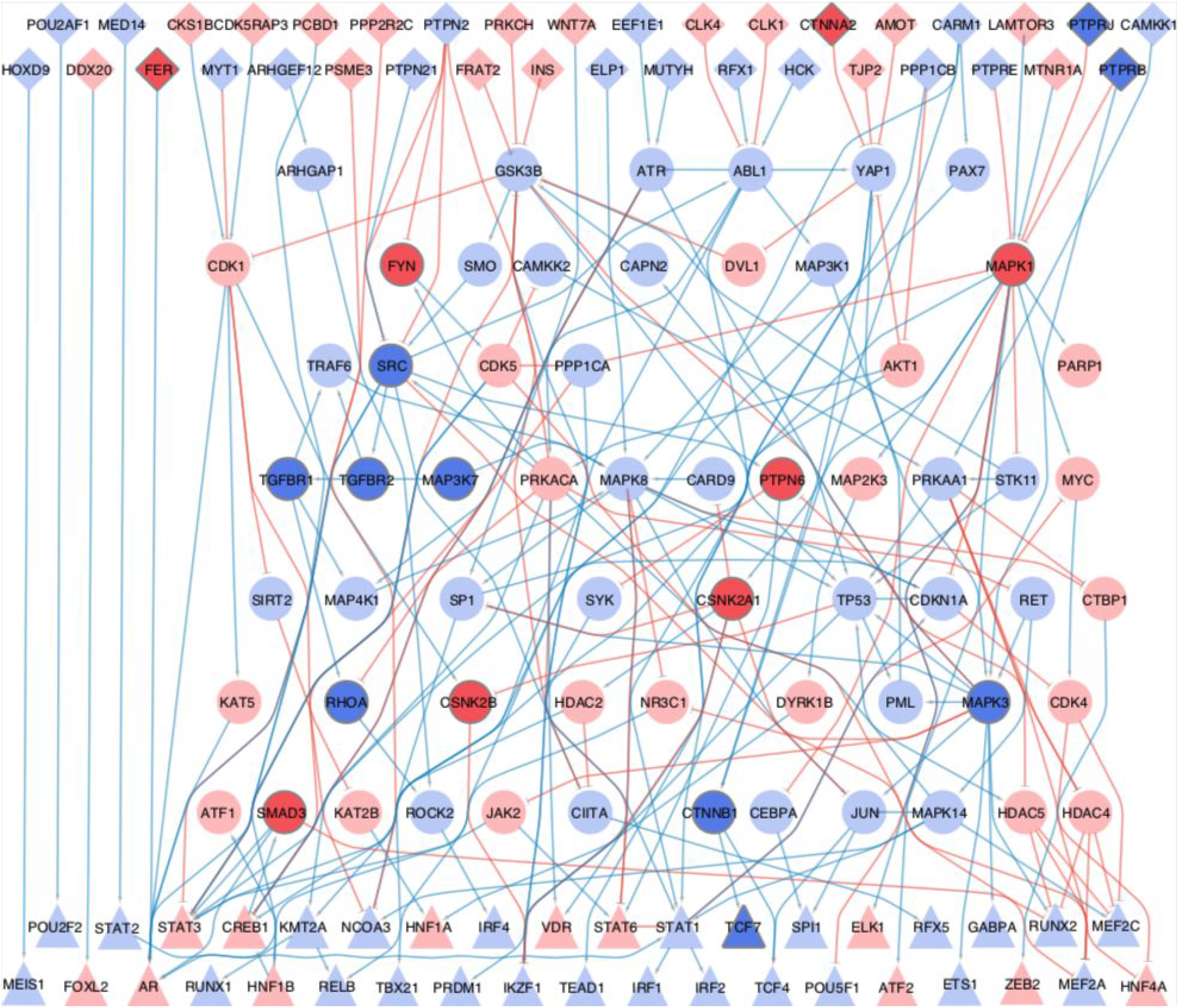
IgAN-contextualized network from CARNIVAL. The network summarizes the CARNIVAL results for node penalty β=0.5. This network consists of 43 TFs, 37 input nodes and 62 associated nodes which are connected through 231 edges. Up-regulated nodes and activatory reactions are indicated in blue while down-regulated nodes and inhibitory edges are colored in red. Triangles correspond to transcription factors, diamonds represent input nodes and circles correspond to purely inferred nodes. Members of the most dysregulated gene set, i.e. adherens junctions, are labeled by more intense background colors..

In addition, we performed down-sampling and network randomization via re-shuffling of TF scores and labels to assess the robustness of CARNIVAL results for the IgAN datasets (see Suppl. Figure S5). We observed that the inferred node activities of CARNIVAL networks from 70% down-sampling datasets is relatively similar to the true IgAN network inferred from the full dataset (Jaccard similarity measure: 0.448 +/− 0.051 [mean +/− S.D.]). In contrast, a major difference was observed when compared the true IgAN network with the randomized networks from the re-shuffling dataset (Jaccard similarity measure: 0.065 +/− 0.032 [mean +/− S.D.]). These results suggest that CARNIVAL provides robust results significantly different from random networks (p-value < 0.001).

#### 3.2.2 Topological analyses and text-mining results of CARNIVAL networks

To identify the central regulatory nodes in the IgAN inferred networks, network’s properties of the results generated from different node penalties parameters were extracted. These measures include in-, out-, and all-degree of interactions as well as betweenness, hub-score and authority-score of nodes in the networks (see Methods). The Top-20 results in each category are shown in Suppl. Table S3 and the robustness of results are shown in Suppl. Figure S6. Of note, signaling molecules which are related to p53 pathway and cell cycle regulation including CDK1, ATM, and TP53 appeared frequently in the top list. In addition, the ERK/MAPK pathway represented by MAPK1 (ERK-2) and MAPK3 (ERK-1) was shown to have high-degree of connectivity where MAPK3 has a very high hub-score. The distributions of network topology measures for these nodes from the IgAN networks are statistically different comparing to the ones from randomized PKNs with the same degree distribution (average p-value = 0.0135). This excludes that these results are because these proteins are hub nodes in the prior knowledge network (PKN), In addition, TP53 is the highest scoring node in randomized networks while MAPK1 amd MAPK3 are often found to rank higher in the IgAN network. Note that a similar finding was observed for GSK-3β (GSK3B), highlighting the potential involvement of PI3K/Akt pathway (together with AKT1) as well as Wnt pathway in the molecular pathogenesis of IgAN.

Besides topological analyses, we systematically searched in the literature to identify whether the signalling proteins identified to be dysregulated in CARNIVAL have support for the roles in the pathophysiology of IgAN (see Methods and Suppl. Table S4). We observed that the molecules in the MAPK pathway and PI3K/Akt pathways, which are previously supported by topological analysis, also have multiple hits (7 hits for MAPK1/MAPK3 and 3 hits for AKT1). On the other hand, we identified many molecules which are in the top list of network topology measures but with no hit on text-mining results. These include down-regulated MAPK14 in the p38 MAPK pathway, down-regulated GSK3B in the PI3K/Akt and Wnt pathways, and up-regulated CDK5 in the cell cycle circuit. Also, we identified RhoA (RHOA) and β-Catenin (CTNNB1) as the components in the TGF-β and Wnt pathways having a few literature support with 2 and 3 hits, respectively. These less or not characterized proteins are candidate players in IgAN, pending validation.

#### 3.2.3 Inferred dysregulated cellular processes by CARNIVAL

Dysregulated cellular processes were inferred by over-representation analysis of the CARNIVAL nodes in the KEGG pathway gene sets. The most significantly dysregulated pathways were identified by median p-value over different node penalties (Figure 5). Among these, known drivers of renal fibrosis including TGF-β, WNT, and EGFR/ERbB signaling stand out (33–35). These processes are significantly over-represented in CARNIVAL, but are not significant in GSEA (results not shown). Additionally, focal adhesions were reported as an activated process in CARNIVAL and GSEA. Interactions between extracellular matrix and the cytoskeleton are particularly important in matrix-producing cells like fibroblasts and mesangial cells in the kidney (36). The enrichments of MAPK, Wnt, TGF-β and cell cycle pathways were also supported by topological network analyses and text-mining results.

**Figure 5.**
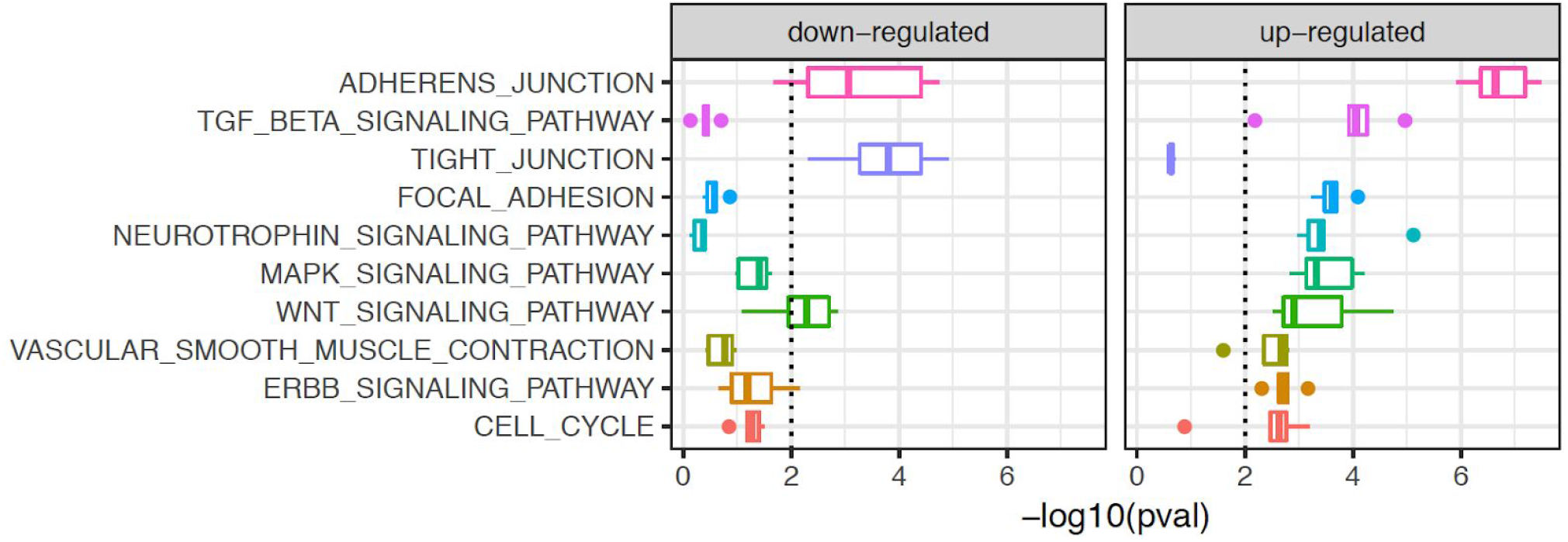
Dysregulated cellular processes in IgAN. Up- and down-regulated pathways are shown with decreasing median significance from top to bottom. The significance level is 0.01. Among others, these point to podocyte injury and the disruption of the slit diaphragms, as well as fibrosis.

Wnt/β-catenin signaling has a central role in mediating podocyte dysfunction and β-catenin is activated in podocytes in various proteinuric kidney diseases, including IgAN (34). Consistent with clinical data in IgA nephropathy reporting podocyte injury and podocyturia as important prognostic features, we identified adhesion junctions as the most significantly dysregulated gene set and tight junctions as the most significantly down-regulated one (37).

While CARNIVAL predicts an upregulation for Wnt and TGF-β (p-value: 0.0012 and 5.47e-5, respectively), GSEA analysis predicts that these two pathways are down-regulated in GSEA (p-values for the down-regulation of pathways in GSEA are 0.0778 and 0.0085, respectively). These pathways are therefore worth to be investigated experimentally to confirm the validity of results from the two approaches.

#### 3.2.4 Fluorescence immunohistology detection of RhoA and β-catenin

The role of MAPK and PI3K/Akt signalling on IgAN’s pathophysiology have been described in previous studies (38–41). We therefore focussed on validating key components of the TGF-β and Wnt signaling pathways with less literature support and where CARNIVAL and GSEA provided opposite results. We chose RhoA (RHOA) and β-catenin (CTNNB1) for fluorescence immunostaining on human renal biopsies from healthy pre-transplantation (controls) and biopsies from diagnosed IgAN patients (see Figure 6 and Suppl. Text S4). Both RhoA and β-catenin genes were down-regulated, as were also the corresponding pathway enrichment scores. In contrast, CARNIVAL predicts that signalling activity is increased for both. Using immunohistology, mesangial IgA (green) was present in IgAN but absent in control specimens as expected for this condition (Figure 6D-F and Figure 6A-C, respectively). RhoA (blue) and β-catenin (red) were present, albeit differentially expressed between IgAN and control. RhoA (blue) was expressed ubiquitously in both kidney’s glomeruli and renal tubules. In comparison to healthy biopsies, we observed an increase in β-catenin staining in IgAN glomeruli, most likely in mesangial cells (Figure 6). Thus, the increase in β-catenin expression might be related to increased Wnt/β-catenin signalling in IgAN glomeruli and this might be related to the deposition of IgA.

**Figure 6.**
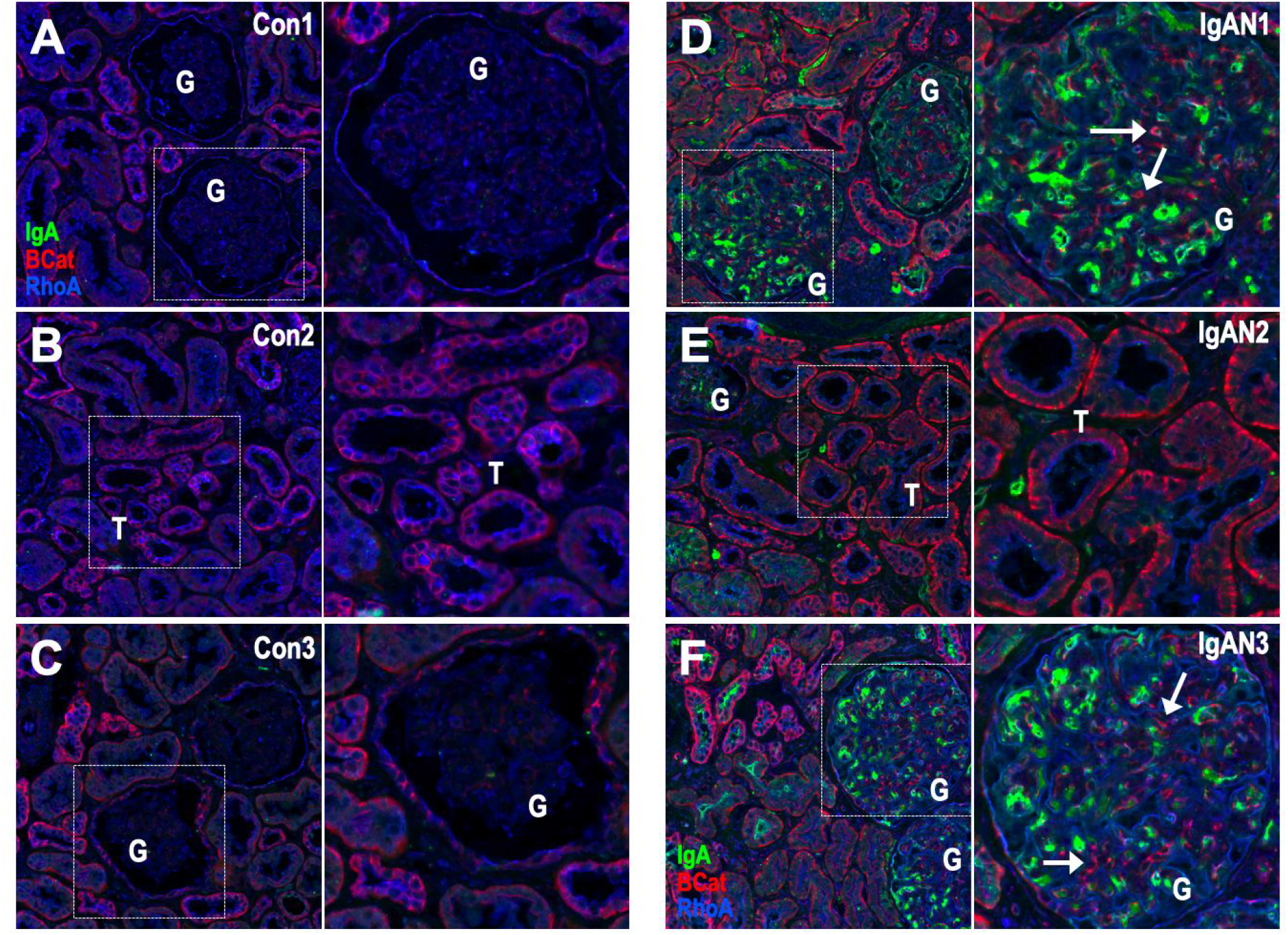
Validation experiment. IgA (green) beta-catenin (red) and RhoA (blue) staining was performed in human biopsies collected from either healthy pre-transplantation control donors (Con1-3; A-C) or diagnosed IgAN patients (IgAN1-3; D-F). 3 representative examples are shown and areas with glomeruli (G) and proximal tubules (T) are indicated. Accumulation of IgA (green) is the pathological hallmark of IgAN and there is no IgA staining in control specimens (A-C). RhoA immunostaining (blue) seems to be ubiquitous and dispersed in tubules and glomeruli. beta-catenin (red) is elevated in IgAN biopsies and there is an increase in beta-catenin cellular staining in glomeruli (arrows). Dotted white boxes depict highlighted areas magnified on left panels. All sections were 10um thick scanned with a 20x lens.

## 4 Discussion

In this paper we present an open-source novel causal network contextualization tool, CARNIVAL. It seamingly integrates gene expression data with various types of prior biological knowledge (signaling networks, TF-targets, and pathway-footprints) to identify processes driving changes in gene expression. The network inference process is swiftly performed with an ILP formulation of causal reasoning. Importantly, using the variant InvCARNIVAL, the origin of these changes (e.g. perturbation targets) do not need to be known to produce the contextualized networks.

According to our benchmarking study, the introduction of TF weights from DoRothEA improves the performance of StdCARNIVAL (Suppl. Figure S1). For InvCARNIVAL, a performance was improved with the additional introduction of pathway weights inferred from PROGENy (Suppl. Figure S2), although the same advantage was not observed in StdCARNIVAL. This demonstrates that the pathway weights only guide the network search if a direction is not provided through a known perturbed target node.

The benchmarking results show that the InvCARNIVAL implementation with TF and pathway weights can obtain perturbation-attributed pathways with a comparable accuracy to StdCARNIVAL (Figure 3). Therefore, we recommend to include the TF and pathway implementations as the default setting. Comparing to GSEA, perturbation-attributed pathways were more frequently identified with the correct direction. Additionally, a related method, CausalR, did not perform as well as InvCARNIVAL or GSEA in detecting up-regulated pathway sets and was biased towards hub nodes while CARNIVAL was not (Suppl. Figure S4).

In the application study in IgAN, besides standard enrichment analyses, quantitative measures from network topology analyses also pointed to the involvement of multiple signalling pathways in the pathophysiology of the disease. These include MAPK, PI3K/Akt, cell cycle, Wnt and TGF-β pathways where we chose the latter two which have less literature support and have conflicting results to GSEA for experimental validation. WNT signaling was reported as a dysregulated process in CARNIVAL and is known to be involved in podocyte injury and renal fibrosis (34). The IgAN network included representative mediators of the classical WNT signaling pathway from the messenger WNT7A to the TFs TCF4 and TCF7, although it should be noted that not all of these are linked in the expected ways nor do all members show the expected activity. GSK3B, which is one of the components in the WNT signaling pathway, also appears among the nodes with top network topology measure, highlighting the importance of this molecule as an important mediator in the signalling regulation of IgAN’s pathophysiology.

TGF-β signaling is a main driver of fibrosis (33). CARNIVAL’s IgAN network captured all members of the TGF-β/RhoA pathway as up-regulated and linked through the biologically expected interactions. This includes the TGF-β receptors (TGFBR1 and TGFBR2), the ras homolog gene family member A (RHOA) and the Rho-associated protein kinase 2 (ROCK2). This is consistent with the previously reported up-regulation of protein levels of TGF-β receptors and RhoA in IgAN (42, 43) and illustrates how CARNIVAL can identify highly relevant and specific processes and regulators from gene expression data. Our validation experiment shows that both β-catenin and RhoA are present in epithelial cells in IgAN, consistent with the involvement of these cells in the pathophysiology of the disease. This was captured neither via differential expression analysis at the individual gene level nor via GSEA at the pathway scale (Figure 6, (32)). β-catenin and Wnt signalling are studied as drug targets for different cancers (44). They have been proposed as a potential target for chronic kidney (45). There are multiple Wnt signalling small-molecule drugs that bind β-catenin (46). Such drugs have not been tested in IgAN thus far. Given the pathological importance of the glomerulus in IgAN, the possible differential expression and signalling function of RhoA and Wnt/β-catenin needs to be investigated further. Other dysregulated signalling molecules identified by CARNIVAL, including MAPK14, GSK3B and CDK5, are also interesting targets for further validation, as their role on the pathophysiology of IgAN is unknown.

Overall, we demonstrate the superior performance of CARNIVAL over existing methods in the benchmarking study and also its applicability to biomedical data in our IgAN case study. However, it should be noted that the benchmarking is performed at the cellular process level due to the limited information on protein activities (see Methods). Moreover, the two-step inference approach also has a few limitations: 1) Not all attributed pathways in the KEGG database are represented in MSigDB nor equally relevant, 2) Gene sets do not account for directionality and 3) Gene sets for the same process can be inconsistent in different databases. All of these factors could affect the benchmarking results across all methodologies being tested, and further analyses should be performed to determine the generality of our findings.

Although we demonstrated that incorporating prior knowledge into the network inference can lead to a higher accuracy, the drawback is the inherent bias towards known biology. CARNIVAL only uses the known interactions as a scaffold and the contextualization of the network is data-driven. Hence, it can still be applied to predict the status of proteins and their connections for specific contexts. Since CARNIVAL can not propose *de novo* connections between signaling molecules, it could be combined with pure data-driven network inference approaches such as nested effect models (NEM) (47) in the future.

To conclude, we believe that, given the flexibility of the CARNIVAL pipeline, it can be a useful tool to infer context-specific signaling network architectures from gene expression in many studies.

## 5 Acknowledgements

This work was supported by the European Union’s Horizon 2020 program [675585 Marie-Curie ITN “SymBioSys”]; and the Innovative Medicines Initiative 2 Joint Undertaking [116030 “TransQST”]. This Joint Undertaking receives support from the European Union’s Horizon 2020 research and innovation programme and EFPIA.

We also thank Aurélien Dugourd for sharing the *runPROGENy* script and for the inputs for benchmarking using enrichment analysis, Bence Szalai, Luz Garcia-Alonso and Luis Tobalina for feedback on the CARNIVAL pipeline and Attila Gábor for the support on compiling the CARNIVAL package.

## 6 Authors contributions

AL performed benchmarking and IgAN studies. PT compiled the CARNIVAL workflow as an R-package. EG implemented the ILP formulation in R. AL, PT and EG designed the CARNIVAL pipeline, analysed results and wrote manuscript. AD and JB analysed IgAN results and performed the validation experiment. JSR conceived the project. PT and JSR supervised the project. All authors read and revised the manuscript.

