## Supplementary Information for "From expression footprints to causal pathways: contextualizing large signaling networks with CARNIVAL"

**Supplementary Figures:**

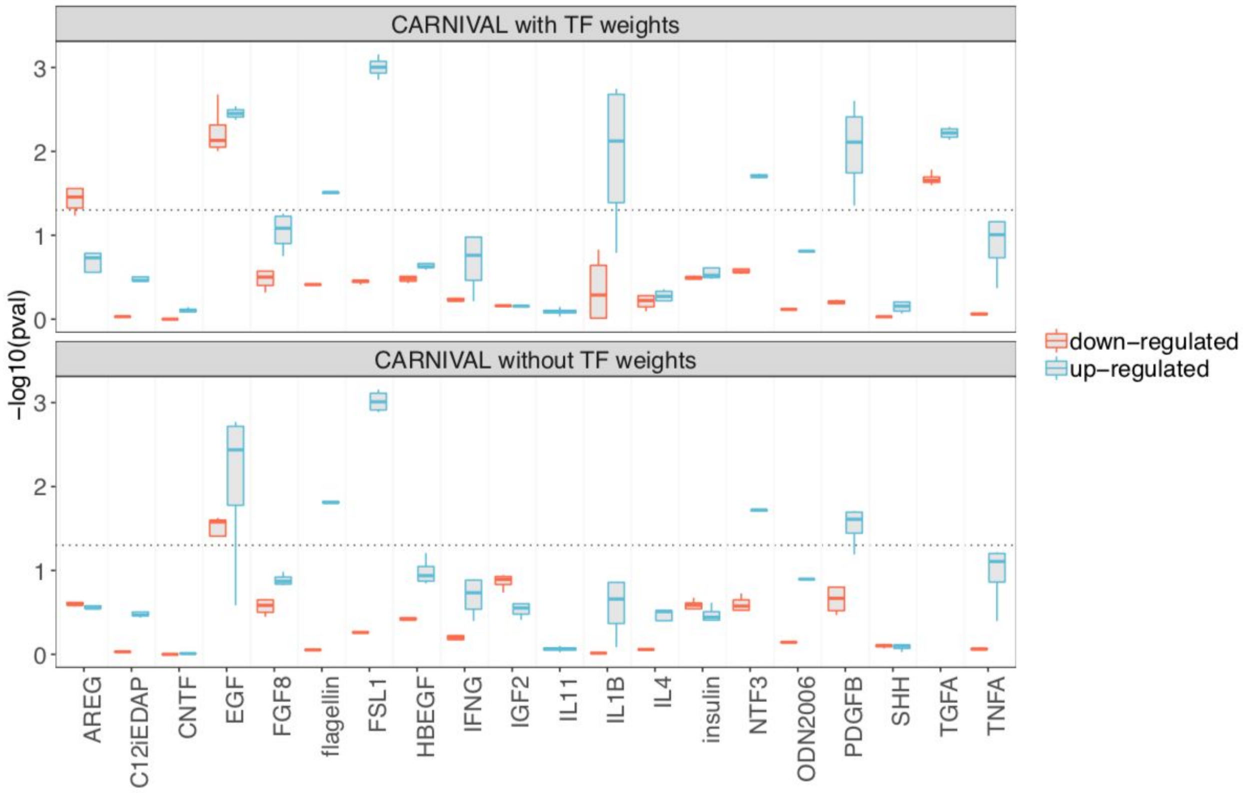


**Figure S1**. Results from with and without the integration of TF weights in standard CARNIVAL. Results are shown for beta = 1.

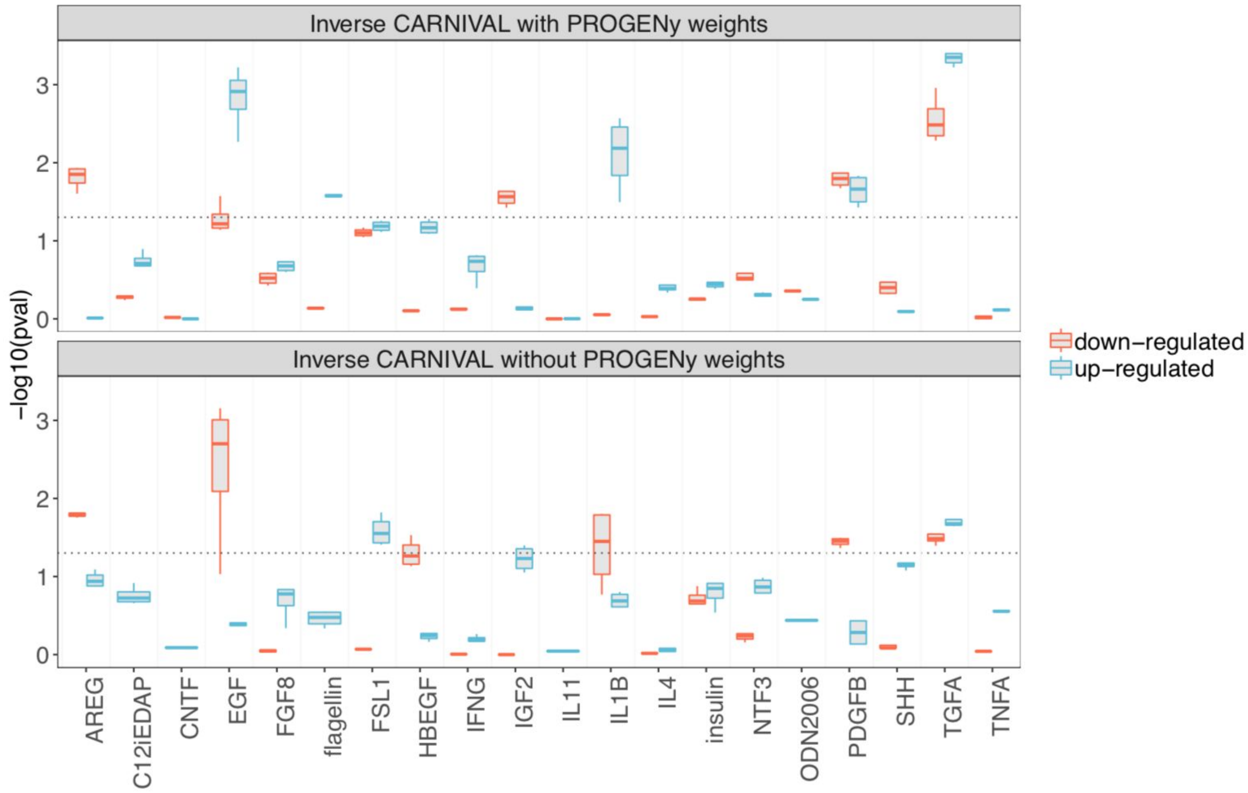


**Figure S2**. Results from the integration of pathway weights in inverse CARNIVAL. Results are shown for beta = 0.5.

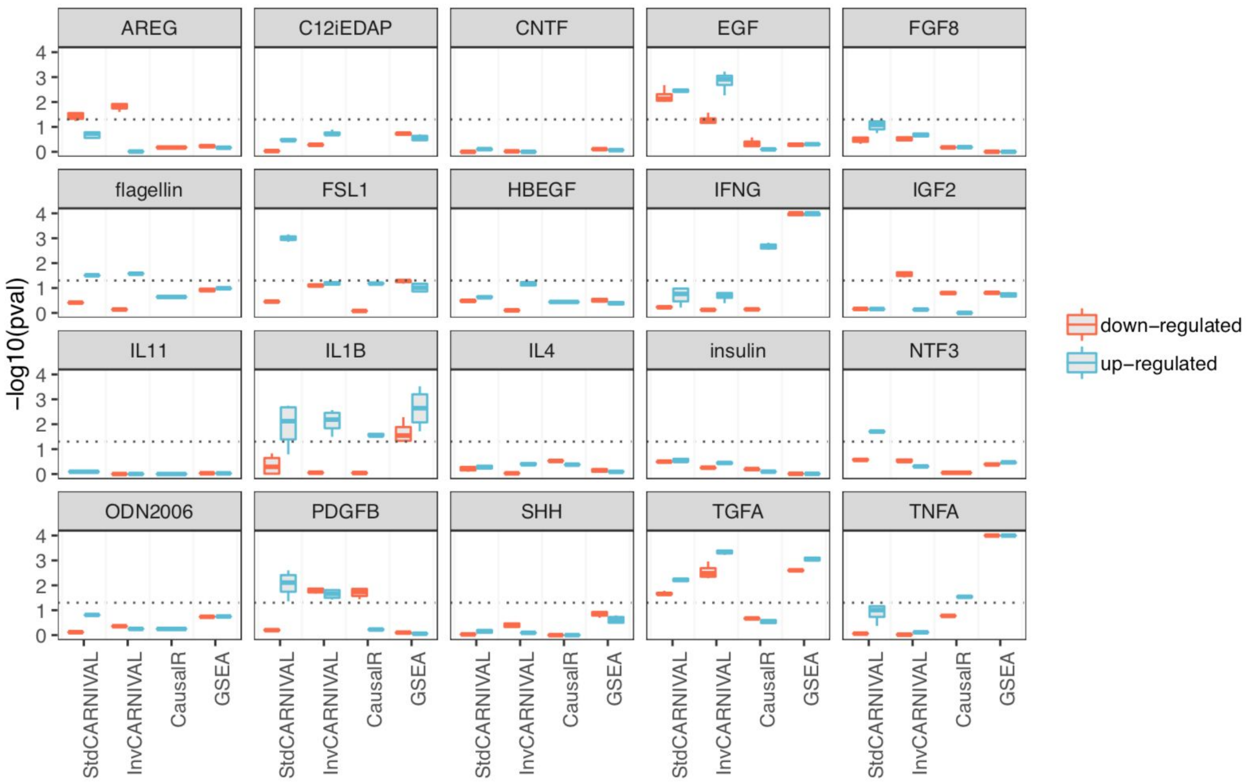


**Figure S3**. Comparison of the enrichment results of the perturbation-attributed pathway set in dysregulated pathways inferred with different tools. An enrichment of the perturbation-attributed pathway set among the significant pathways was determined. For GSEA, dysregulated pathways were inferred with the *piano* R-package to determine whether a certain pathway is activated (up-regulated) or inhibited (down-regulated). For CausalR and CARNIVAL, an over-representation analysis of up- or down-regulated solution nodes in the KEGG gene sets was used as a proxy instead. The significance level of 0.05 is indicated by the dotted lines.

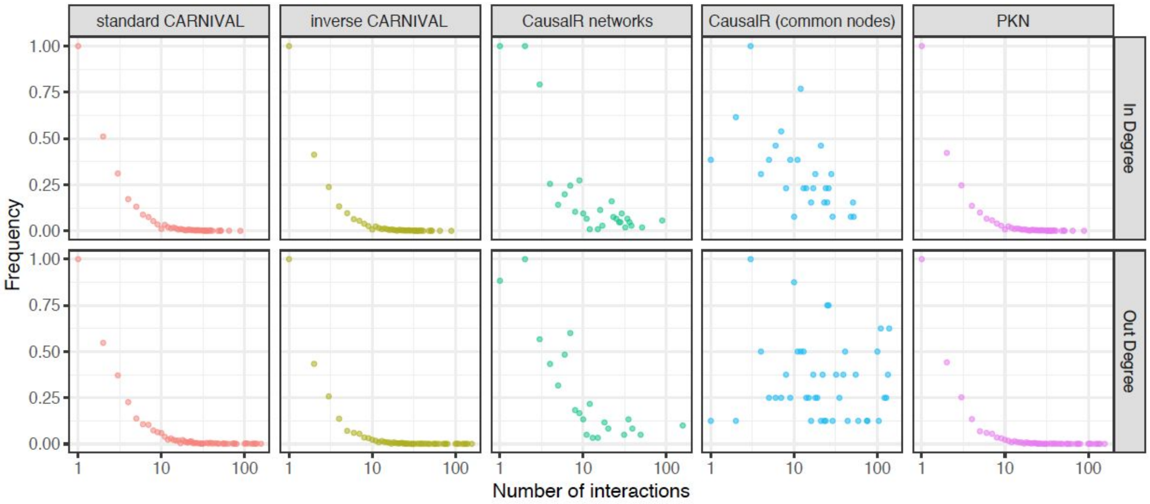


**Figure S4**. Comparison of node connectivity distributions for networks from CARNIVAL and CausalR comparing to the prior knowledge network (PKN).

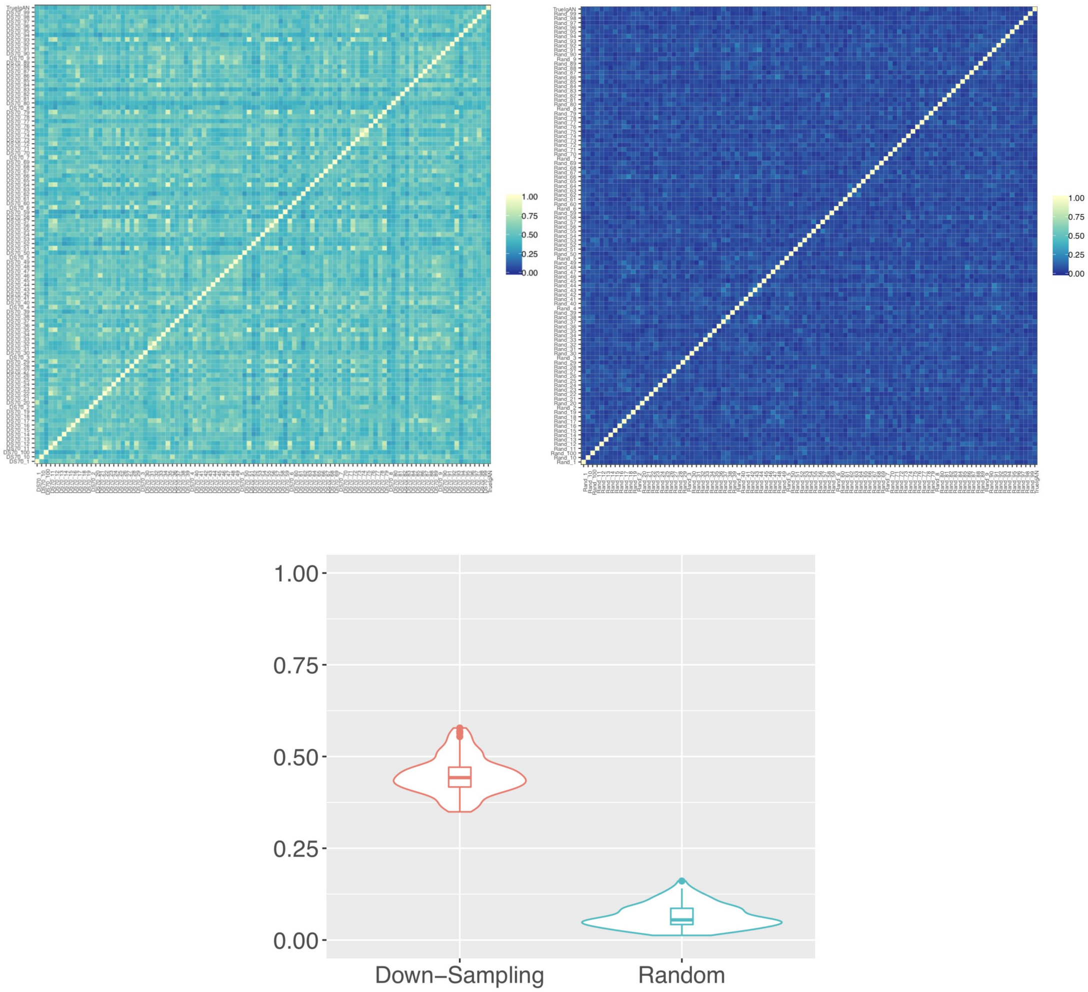


**Figure S5**. Jaccard similarity measures of 100 networks generated by 70% down-sampling (top-left) and by re-shuffling of transcription factor (TF) inputs' labels (top-right) comparing to the true IgAN network from actual data. The mean and standard deviation of the Jaccard similarity measure of the 70% down-sampling networks is 0.448 +/- 0.051 (mean +/- S.D.) while they are 0.065 +/- 0.032 (mean +/- S.D.) from TF-labels reshuffling (bottom). The p-value from one-sided Komogorov-Smirnov test between the two sets of Jaccard similarity distribution is 3.72e-44.

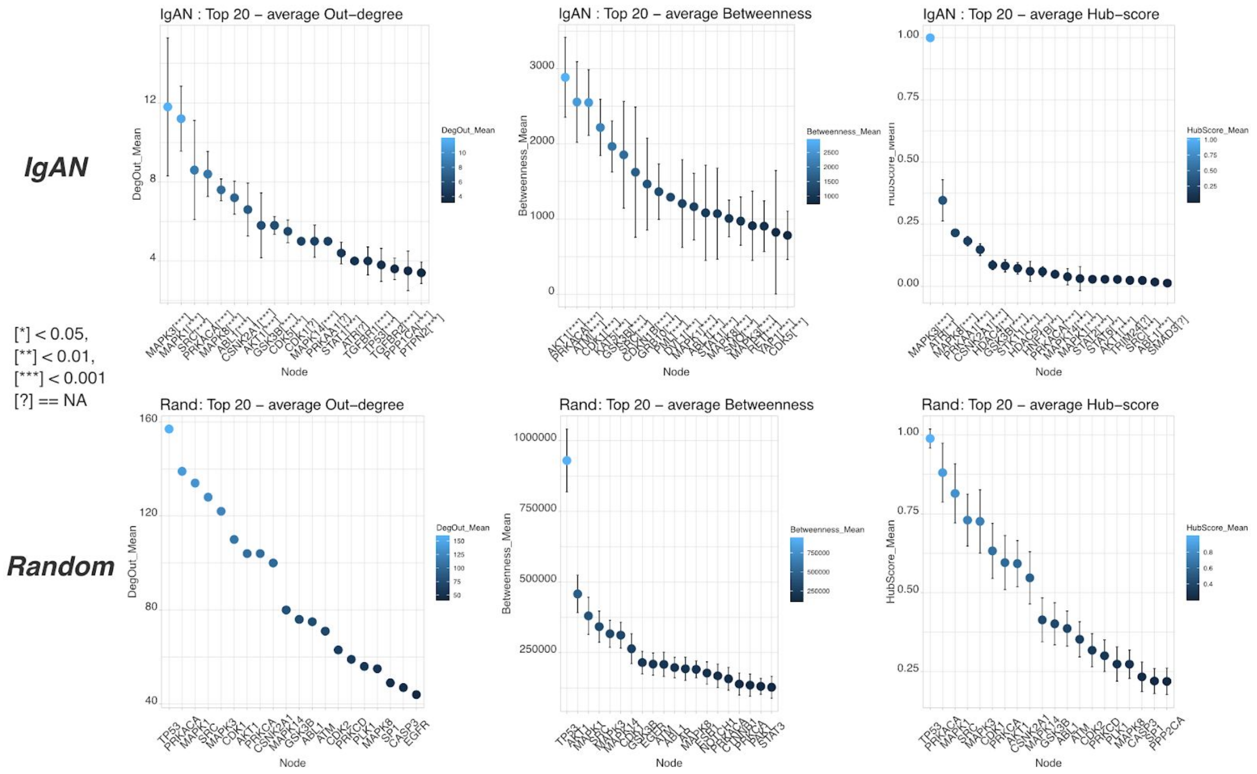


**Figure S6**. Network topology measures from IgAN-contextualized networks from CARNIVAL at five different size penalty (beta=0.03,0.1,0.3,0.5,0.8) and from 100 randomized networks with degree and sign preserved generated from the prior knowledge network. Dots and error bars represent the mean and standard deviation for each node in each category. Significant differences from t-test are highlighted with asterisks (*) after the protein names in the IgAN set. Question marks (?) were annotated when a significant test could not performed e.g. due to limited sample sizes.

**Supplementary Tables:**

**Table S1**. Representative nodes of PROGENy pathway in CARNIVAL. A minimal number of representative nodes were chosen in a way that covers all associated signalling routes described in KEGG while avoiding overlaps between pathways and with DoRothEA TFs if possible.

| Pathway | Representative Nodes |
|---|---|
| EGFR | EGFR, ERBB2 |
| Hypoxia | HIF1A |
| JAK/STAT | JAK1, JAK2, JAK3 |
| MAPK | BRAF, ARAF, RAF1 |
| NFκB | NFKB1 |
| PI3K | PIK3CA, PIK3CB, PIK3CD, PIK3CG |
| TGFβ | TGFBR1, TGFBR2, BMPR1A, BMPR1B, BMPR2 |
| TNFα | TNFRSF1A, TNFRSF1B |
| Trail | CASP8, CASP10 |
| VEGF | FLT1, FLT3, KDR, PDGFRA, PDGFRB |
| p53 | TP53 |
| Androgen | AR |
| Estrogen | ESR1, ESR2 |
| WNT | DVL1 |

**Table S2**. Glomerular gene expression datasets. Microarray gene expression data for IgAN patients and healthy living donors (HLD) were accessed from the GEO database (1). The accession number is shown together with the number of samples in each class (IgAN, HLD).

| Study | GEO accession | Platform | HLD | IgAN |
|---|---|---|---|---|
| Berthier et al. (2012) (2) | GSE37460 | GPL96 & GPL570 | 27 | 27 |
| Hodgin et al. (2014) (3) | GSE50469 | GPL96 & GPL570 | 0 | 22 |
| Liu et al. (2017) (4) | GSE93789 | GPL570 | 22 | 20 |
| Woroniecka et al. (2011) (5) | GSE30122 | GPL571 | 6 | 0 |
| Berthier et al. (2012) (2) | GSE32591 | GPL96 | 4 | 0 |

**Table S3**. Top 20 nodes and edges with highest average network topology measures from the IgAN networks with different node penalty parameters (β in (0.03; 0.1; 0.3; 0.5; 0.8), see Methods).

| Top 20 In-degree | | | | | | |
|---|---|---|---|---|---|---|
| Protein | In-degree | Out-degree | All-degree | Betweenness | Hub-score | Authority-score |
| AR | **8.2** | 0 | 8.2 | 0 | 0.0000 | 0.0095 |
| TP53 | **7.4** | 3.8 | 11.2 | 609.2 | 0.0024 | 1.0000 |
| STAT3 | **7.2** | 0.4 | 7.6 | 135.3 | 0.0000 | 0.0302 |
| MAPK1 | **6.4** | 11.2 | 17.6 | 1164.2 | 0.0308 | 0.0002 |
| STAT1 | **6.4** | 4.4 | 10.8 | 1071.7 | 0.0001 | 0.1163 |
| MAPK8 | **5.6** | 7.6 | 13.2 | 1007.1 | 0.2150 | 0.0000 |
| ATM | **5.2** | 3 | 8.2 | 2550 | 0.0027 | 0.0002 |
| GSK3B | **5.2** | 5.8 | 11 | 1854.6 | 0.0821 | 0.0142 |
| CDKN1A | **5** | 1 | 6 | 170.125 | 0.0000 | 0.0056 |
| CREB1 | **5** | 0.2 | 5.2 | 22 | 0.0000 | 0.2396 |
| MEF2A | **5** | 0 | 5 | 0 | 0.0000 | 0.0453 |
| CDK1 | **4.8** | 5 | 9.8 | 2218.3 | 0.0016 | 0.0040 |
| MEF2C | **4.8** | 0 | 4.8 | 0 | 0.0000 | 0.0492 |
| SP1 | **4.8** | 1.8 | 6.6 | 76.4 | 0.0001 | 0.9537 |
| ABL1 | **4.6** | 7.2 | 11.8 | 1082.8 | 0.0170 | 0.0060 |
| IKZF1 | **4.6** | 0 | 4.6 | 0 | 0.0000 | 0.0522 |
| SRC | **4.4** | 8.6 | 13 | 1623.6 | 0.0234 | 0.0020 |
| STAT6 | **4.2** | 1 | 5.2 | 137.6 | 0.0282 | 0.0589 |
| YAP1 | **4.2** | 2.6 | 6.8 | 824.1 | 0.0023 | 0.0137 |
| CTNNB1 | **3.8** | 2 | 5.8 | 86.2 | 0.0000 | 0.0083 |
| Top 20 Out-degree | | | | | | |
| Protein | In-degree | Out-degree | All-degree | Betweenness | Hub-score | Authority-score |
| MAPK3 | 3 | **11.8** | 14.8 | 910.5 | 1.0000 | 0.0000 |
| MAPK1 | 6.4 | **11.2** | 17.6 | 1164.2 | 0.0308 | 0.0002 |
| SRC | 4.4 | **8.6** | 13 | 1623.6 | 0.0234 | 0.0020 |
| PRKACA | 2 | **8.4** | 10.4 | 2557.4 | 0.0483 | 0.0079 |
| MAPK8 | 5.6 | **7.6** | 13.2 | 1007.1 | 0.2150 | 0.0000 |
| ABL1 | 4.6 | **7.2** | 11.8 | 1082.8 | 0.0170 | 0.0060 |
| CSNK2A1 | 1 | **6.6** | 7.6 | 195.4 | 0.1474 | 0.0124 |
| AKT1 | 3.4 | **5.8** | 9.2 | 2885.6 | 0.0277 | 0.0057 |
| GSK3B | 5.2 | **5.8** | 11 | 1854.6 | 0.0821 | 0.0142 |
| CDK5 | 1 | **5.5** | 6.5 | 783.5 | 0.0014 | 0.0000 |
| CDK1 | 4.8 | **5** | 9.8 | 2218.3 | 0.0016 | 0.0040 |
| MAPK14 | 1 | **5** | 6 | 263.125 | 0.0382 | 0.0000 |
| PRKAA1 | 2.4 | **5** | 7.4 | 114.4 | 0.1819 | 0.0160 |
| STAT1 | 6.4 | **4.4** | 10.8 | 1071.7 | 0.0001 | 0.1163 |
| ATR | 2.8 | **4** | 6.8 | 324 | 0.3459 | 0.0059 |

| | | | | | | |
|---|---|---|---|---|---|---|
| TGFBR1 | 1.6 | **4** | 5.6 | 68.8 | 0.0018 | 0.0000 |
| TP53 | 7.4 | **3.8** | 11.2 | 609.2 | 0.0024 | 1.0000 |
| TGFBR2 | 1.6 | **3.6** | 5.2 | 337 | 0.0000 | 0.0068 |
| PPP1CA | 1.25 | **3.5** | 4.75 | 631.25 | 0.0021 | 0.0002 |
| PTPN2 | 0 | **3.4** | 3.4 | 0 | 0.0037 | 0.0000 |
| Top 20 All-degree | | | | | | |
| Protein | In-degree | Out-degree | All-degree | Betweenness | Hub-score | Authority-score |
| MAPK1 | 6.4 | 11.2 | **17.6** | 1164.2 | 0.0308 | 0.0002 |
| MAPK3 | 3 | 11.8 | **14.8** | 910.5 | 1.0000 | 0.0000 |
| MAPK8 | 5.6 | 7.6 | **13.2** | 1007.1 | 0.2150 | 0.0000 |
| SRC | 4.4 | 8.6 | **13** | 1623.6 | 0.0234 | 0.0020 |
| ABL1 | 4.6 | 7.2 | **11.8** | 1082.8 | 0.0170 | 0.0060 |
| TP53 | 7.4 | 3.8 | **11.2** | 609.2 | 0.0024 | 1.0000 |
| GSK3B | 5.2 | 5.8 | **11** | 1854.6 | 0.0821 | 0.0142 |
| STAT1 | 6.4 | 4.4 | **10.8** | 1071.7 | 0.0001 | 0.1163 |
| PRKACA | 2 | 8.4 | **10.4** | 2557.4 | 0.0483 | 0.0079 |
| CDK1 | 4.8 | 5 | **9.8** | 2218.3 | 0.0016 | 0.0040 |
| AKT1 | 3.4 | 5.8 | **9.2** | 2885.6 | 0.0277 | 0.0057 |
| AR | 8.2 | 0 | **8.2** | 0 | 0.0000 | 0.0095 |
| ATM | 5.2 | 3 | **8.2** | 2550 | 0.0027 | 0.0002 |
| CSNK2A1 | 1 | 6.6 | **7.6** | 195.4 | 0.1474 | 0.0124 |
| STAT3 | 7.2 | 0.4 | **7.6** | 135.3 | 0.0000 | 0.0302 |
| PRKAA1 | 2.4 | 5 | **7.4** | 114.4 | 0.1819 | 0.0160 |
| ATR | 2.8 | 4 | **6.8** | 324 | 0.3459 | 0.0059 |
| YAP1 | 4.2 | 2.6 | **6.8** | 824.1 | 0.0023 | 0.0137 |
| HDAC5 | 3.6 | 3 | **6.6** | 36.6 | 0.0604 | 0.0379 |
| SP1 | 4.8 | 1.8 | **6.6** | 76.4 | 0.0001 | 0.9537 |
| Top 20 Betweenness | | | | | | |
| Protein | In-degree | Out-degree | All-degree | Betweenness | Hub-score | Authority-score |
| AKT1 | 3.4 | 5.8 | 9.2 | **2885.6** | 0.0277 | 0.0057 |
| PRKACA | 2 | 8.4 | 10.4 | **2557.4** | 0.0483 | 0.0079 |
| ATM | 5.2 | 3 | 8.2 | **2550** | 0.0027 | 0.0002 |
| CDK1 | 4.8 | 5 | 9.8 | **2218.3** | 0.0016 | 0.0040 |
| KAT5 | 1.4 | 2 | 3.4 | **1966.5** | 0.0003 | 0.0001 |
| GSK3B | 5.2 | 5.8 | 11 | **1854.6** | 0.0821 | 0.0142 |
| SRC | 4.4 | 8.6 | 13 | **1623.6** | 0.0234 | 0.0020 |
| CDKN1B | 2 | 1 | 3 | **1466** | 0.0001 | 0.0000 |
| GRB10 | 2 | 1 | 3 | **1364** | 0.0000 | 0.0159 |
| PML | 2 | 1 | 3 | **1291.25** | 0.0001 | 0.0252 |
| DVL1 | 1 | 1 | 2 | **1206.166667** | 0.0000 | 0.0000 |
| MAPK1 | 6.4 | 11.2 | 17.6 | **1164.2** | 0.0308 | 0.0002 |
| ABL1 | 4.6 | 7.2 | 11.8 | **1082.8** | 0.0170 | 0.0060 |
| STAT1 | 6.4 | 4.4 | 10.8 | **1071.7** | 0.0001 | 0.1163 |

| | | | | | | |
|---|---|---|---|---|---|---|
| MAPK8 | 5.6 | 7.6 | 13.2 | **1007.1** | 0.2150 | 0.0000 |
| SMO | 1 | 1 | 2 | **972.625** | 0.0001 | 0.0054 |
| MAPK3 | 3 | 11.8 | 14.8 | **910.5** | 1.0000 | 0.0000 |
| RET | 1 | 2 | 3 | **905.125** | 0.0000 | 0.0025 |
| YAP1 | 4.2 | 2.6 | 6.8 | **824.1** | 0.0023 | 0.0137 |
| CDK5 | 1 | 5.5 | 6.5 | **783.5** | 0.0014 | 0.0000 |
| Top 20 Hub-score | | | | | | |
| Protein | In-degree | Out-degree | All-degree | Betweenness | Hub-score | Authority-score |
| MAPK3 | 3 | 11.8 | 14.8 | 910.5 | **1.0000** | 0.0000 |
| ATR | 2.8 | 4 | 6.8 | 324 | **0.3459** | 0.0059 |
| MAPK8 | 5.6 | 7.6 | 13.2 | 1007.1 | **0.2150** | 0.0000 |
| PRKAA1 | 2.4 | 5 | 7.4 | 114.4 | **0.1819** | 0.0160 |
| CSNK2A1 | 1 | 6.6 | 7.6 | 195.4 | **0.1474** | 0.0124 |
| HDAC4 | 2.8 | 3 | 5.8 | 57.7 | **0.0851** | 0.0497 |
| GSK3B | 5.2 | 5.8 | 11 | 1854.6 | **0.0821** | 0.0142 |
| STK11 | 1 | 3 | 4 | 417.4 | **0.0722** | 0.0067 |
| HDAC5 | 3.6 | 3 | 6.6 | 36.6 | **0.0604** | 0.0379 |
| HNF1B | 1 | 1.8 | 2.8 | 73.2 | **0.0588** | 0.0000 |
| PRKACA | 2 | 8.4 | 10.4 | 2557.4 | **0.0483** | 0.0079 |
| MAPK14 | 1 | 5 | 6 | 263.125 | **0.0382** | 0.0000 |
| MAPK1 | 6.4 | 11.2 | 17.6 | 1164.2 | **0.0308** | 0.0002 |
| STAT2 | 1.4 | 1 | 2.4 | 71.6 | **0.0282** | 0.0001 |
| STAT6 | 4.2 | 1 | 5.2 | 137.6 | **0.0282** | 0.0589 |
| AKT1 | 3.4 | 5.8 | 9.2 | 2885.6 | **0.0277** | 0.0057 |
| TRIM24 | 0 | 2 | 2 | 0 | **0.0239** | 0.0000 |
| SRC | 4.4 | 8.6 | 13 | 1623.6 | **0.0234** | 0.0020 |
| ABL1 | 4.6 | 7.2 | 11.8 | 1082.8 | **0.0170** | 0.0060 |
| SMAD3 | 1 | 3 | 4 | 46 | **0.0126** | 0.0000 |
| Top 20 Authority-score | | | | | | |
| Protein | In-degree | Out-degree | All-degree | Betweenness | Hub-score | Authority-score |
| TP53 | 7.4 | 3.8 | 11.2 | 609.2 | 0.0024 | **1.0000** |
| SP1 | 4.8 | 1.8 | 6.6 | 76.4 | 0.0001 | **0.9537** |
| RUNX2 | 3.4 | 0 | 3.4 | 0 | 0.0000 | **0.7359** |
| GABPA | 1.2 | 0 | 1.2 | 0 | 0.0000 | **0.6918** |
| ETS1 | 1.2 | 0 | 1.2 | 0 | 0.0000 | **0.6910** |
| JUN | 3.8 | 2.2 | 6 | 111.1 | 0.0000 | **0.4111** |
| CREB1 | 5 | 0.2 | 5.2 | 22 | 0.0000 | **0.2396** |
| KMT2A | 2 | 3 | 5 | 380.2 | 0.0001 | **0.1566** |
| PTPN6 | 1.4 | 2.2 | 3.6 | 175.8 | 0.0030 | **0.1449** |
| CTBP1 | 2 | 1 | 3 | 69.4 | 0.0000 | **0.1371** |
| JAK2 | 1 | 2 | 3 | 23.2 | 0.0039 | **0.1284** |
| STAT1 | 6.4 | 4.4 | 10.8 | 1071.7 | 0.0001 | **0.1163** |
| HNF4A | 2 | 0 | 2 | 0 | 0.0000 | **0.0865** |

| | | | | | | |
|---|---|---|---|---|---|---|
| VDR | 2.2 | 0 | 2.2 | 0 | 0.0000 | **0.0689** |
| STAT6 | 4.2 | 1 | 5.2 | 137.6 | 0.0282 | **0.0589** |
| NR3C1 | 2 | 1 | 3 | 18 | 0.0000 | **0.0534** |
| IKZF1 | 4.6 | 0 | 4.6 | 0 | 0.0000 | **0.0522** |
| HDAC4 | 2.8 | 3 | 5.8 | 57.7 | 0.0851 | **0.0497** |
| MEF2C | 4.8 | 0 | 4.8 | 0 | 0.0000 | **0.0492** |
| MEF2A | 5 | 0 | 5 | 0 | 0.0000 | **0.0453** |

**Table S4.** Results from literature search based on the list of up- and down-regulated nodes identified by CARNIVAL. 'Hits' refers to the number of these nodes being present together with the IgAN term. The suffixes '_up' and '_dn' refers to the activity of the nodes in CARNIVAL networks. To account for consistencies, only the nodes which are present in at least 4 out of 5 size penalty parameters (beta) being tested are included. No conflicting CARNIVAL node's activity was observed.

| Hits | CARNIVAL nodes |
|---|---|
| 8 | JUN_up |
| 7 | MAPK1_dn, MAPK3_up, SYK_up |
| 6 | STAT3_dn |
| 4 | CDK1_dn, MAPK8_up |
| 3 | AKT1_dn, CTNNB1_up, MYC_dn |
| 2 | CARD9_up, GABPA_up, JAK2_dn, RHOA_up, SP1_up, SRC_up, STAT1_up, STAT6_dn, VDR_dn, ZEB2_dn |
| 1 | ATF1_dn, CDKN1A_up, CDKN1B_up, HDAC2_dn, HDAC4_dn, KMT2A_up, PRKACA_dn, TBX21_up, TP53_up, TRAF6_up |
| 0 | ABL1_up, ACTL6A_dn, AR_dn, ARHGAP1_up, ARHGEF12_up, ATAD2_dn, ATF2_dn, ATM_dn, ATR_up, CAMKK1_up, CARM1_up, CDK4_dn, CDK5_dn, CDK5RAP3_dn, CDKN2B_up, CEBPA_up, CIITA_up, CKS1B_dn, CLK1_dn, CREB1_dn, CSNK2A1_dn, CSNK2B_dn, CTBP1_dn, DDX20_dn, DUSP7_up, DYRK1B_dn, EEF1E1_up, ELK1_dn, ETS1_up, FOXL2_dn, FYN_dn, GSK3B_up, HCK_up, HDAC5_dn, HNF1A_dn, HNF1B_dn, HNF4A_dn, HOXD9_up, IKZF1_up, IRF1_up, IRF2_up, IRF4_up, KAT2B_dn, KAT5_dn, LAMTOR3_dn, MAP2K3_dn, MAP3K1_up, MAP3K7_up, MAP4K1_up, MAPK14_up, MED14_up, MEF2A_up, MEF2C_up, MEIS1_up, MTNR1A_dn, MUTYH_up, MYT1_up, NCOA3_up, PAX7_up, PCBD1_dn, POU2AF1_up, POU2F2_up, POU5F1_up, PPP1CA_up, PPP1CB_up, PPP2R2C_dn, PRDM1_up, PRKAA1_up, PTPN21_up, PTPN6_dn, PTPRB_up, PTPRJ_up, RAD50_dn, RELB_up, RET_up, RFX1_up, RFX5_up, ROCK2_up, RUNX1_up, RUNX2_up, SIRT2_up, SMO_up, SPI1_up, STAT2_up, STK11_up, TCF4_up, TCF7_up, TEAD1_up, TGFBR1_up, TGFBR2_up, WNT7A_dn, YAP1_up |

**Supplementary Texts:**

**Text S1. ILP implementation.**

We re-implemented the integer linear programming (ILP) formulation of causal reasoning problem presented in Melas et al. (6) in the R programming language. A summarised description of the ILP formation is shown as follows:

A signaling network $G$ is an interaction graph defined by a set of signed and directed reactions $i = 1, 2, ..., n_r$, and a set of nodes $j = 1, 2, ..., n_s$. Thereby, each reaction $i$ is characterized by an ordered pair consisting of source species $S_i$ and target species $T_i$ with $S_i, T_i \in \{1, 2, ..., n_s\}$. The reaction sign is denoted by $\sigma_i \in \{-1, 1\}$ and distinguishes between activation ($\sigma_i = 1$) and inhibition ($\sigma_i = -1$). The ILP variable definitions are summarized in Table 1 and provide a key to the used notation.

**Table 1.** The list of ILP variables and their descriptions

| **Variable** | **Description** |
|---|---|
| $u_i^+ \in \{0, 1\}$ | Potential of reaction $i$ to activate its target node |
| $u_i^- \in \{0, 1\}$ | Potential of reaction $i$ to inhibit its target node |
| $\sigma_i \in \{-1, 1\}$ | Sign of reaction $i$ |
| $x_j^+ \in \{0, 1\}$ | Potential of node $j$ to be activated |
| $x_j^- \in \{0, 1\}$ | Potential of node $j$ to be inhibited |
| $x_j \in \{-1, 0, 1\}$ | Predicted activation/inhibition state of species $j$ |
| $m_j \in \{-1, 0, 1\}$ | Activation/inhibition state of measured species $j$ |
| $I_j \in \{-1, 0, 1, NaN\}$ | Activation/inhibition state of perturbed species $j$; $NaN$ represents the unknown state |
| $B_j \in \{-1, 0, 1\}$ | Auxiliary variable to determine the state perturbed species $j$ once the input value $I_j = NaN$ |
| $d_j \in [0, M]$ | Auxiliary distance variables assigned to each node $j$ where $M$ is a sufficiently large number (default: $M = 100$) |
| $A_j \in [0, 2]$ | Auxiliary variable representing the absolute difference between the inferred and measured species $j$. |

The variables in Table 1 are determined during the linear programming optimisation according to the following set of constraints of causal reasoning principle. The activation state of a reaction $i$ is defined by the activity of its source node $x_{Si}$ and the reaction sign $\sigma_i$. The reaction has a potential to activate its target node ($u_i^+ = 1$), if and only if $\sigma_i \cdot x_{Si} = 1$. This occurs in two cases: either the source node is activated ($x_j = S_i = 1$) and has an activating effect on its target node ($\sigma_i = 1$); or the source node is inhibited ($x_j = S_i = -1$) and has an inhibiting effect ($\sigma_i = -1$). Vice versa, a reaction has the potential to downregulate its target node ($u_i^- = 1$), if and only if $\sigma_i \cdot x_{Si} = -1$.

$$u_i^+ \geq \sigma_i x_j \text{ where } i \in \{1,2,...,n_r\}\,;\, j \in \{1,2,...,n_s\} \quad \ldots\, c_1$$
$$u_i^- \geq -\sigma_i x_j \text{ where } i \in \{1,2,...,n_r\}\,;\, j \in \{1,2,...,n_s\} \quad \ldots\, c_2$$
$$u_i^+ \leq 1 - u_i^- \text{ where } i \in \{1,2,...,n_r\} \quad \ldots\, c_3$$
$$u_i^+ \leq \sigma_i x_j + u_k^- \text{ where } i \in \{1,2,...,n_r\}\,;\, j \in \{1,2,...,n_s\} \quad \ldots\, c_4$$
$$u_i^- \leq -\sigma_i x_j + u_i^+ \text{ where } i \in \{1,2,...,n_r\}\,;\, j \in \{1,2,...,n_s\} \quad \ldots\, c_5$$

For the definition of the activation state of a node $x_j$, two cases can be distinguished: For the non-input nodes, the activity of these nodes is defined by the potentials of incoming reactions. If and only if at least one incoming reaction has the potential to activate ($u_i^+ : T_{i=j} = 1$), the node can have the potential to be activated ($x_j^+ = 0 \vee 1$). Vice versa, a node can only have the potential to be down-regulated ($x_j^- = 0 \vee 1$) if at least one incoming reaction has the potential to inhibit ($u_i^- : T_{i=j} = 1$). A node is then up-regulated ($x_j = 1$), if there is exclusively a potential to be up-regulated ($x_j^+ = 1$), and is down-regulated ($x_j = -1$) if there is exclusively a potential to be down-regulated ($x_j^- = 1$). If none or both of the potentials exist, the node will remain neutral ($x_j = 0$).

$$x_j^+ \leq \sum_{i:T_i=j} u_i^+ \text{ where } i \in \{1,2,...,n_r\}\,;\, j \in \{1,2,...,n_s\} \quad \ldots\, c_6$$

$$x_j^- \leq \sum_{i:T_i=j} u_i^- \text{ where } i \in \{1,2,...,n_r\}\,;\, j \in \{1,2,...,n_s\} \quad \ldots\, c_7$$

For perturbed/input nodes, the activation state can be user-defined ($x_j = I_j$). In the case where the perturbed node is known but the activation/inhibition state remains unknown, the notation $NaN$ can be assigned. The state of $x_j$ is equal to $B_j$ in this case and will then take any of the three states $x_j \in \{-1,0,1\}$. All remaining input nodes which were not defined for its activation state in the input list will always take the value 0 and will not influence the system.

$$x_j = x_j^+ - x_j^- + B_j \text{ where } j \in \{1,2,...,n_s\} \quad \ldots\, c_8\ [1]$$
$$x_j = I_j \text{ where } j \in I\,;\, I_j \in \{-1,0,1\} \quad \ldots\, c_8\ [2]$$

$$x_j = B_j \text{ where } j \in S - T \qquad \ldots\ c_8\ [3]$$
$$B_j = 0 \text{ where } j \in (S \cup T) - I \qquad \ldots\ c_8\ [4]$$

In addition, feedback loops are removed given that effects mediated through those are highly dynamic and hardly interpretable from a static snapshot as in an interaction network. For instance, positive feedback loops can lead to internal signals independent to external perturbations. These were constrained through a distance variable $d_j$. Thereby, the distance of all nodes connected to a perturbation node is set to a value larger than zero, while all others are defined to be zero. As a consequence, only nodes connected to a perturbation can be deregulated. The distance increases from source node to target node if the interaction is active, i.e. $u_i^+ = 1 \vee u_i^- = 1$, and is not allowed to pass the distance threshold $M$, which is considerably larger than expected path lengths.

$$x_i^+ \leq d_j \qquad \ldots\ c_l\ [1]$$
$$x_i^- \leq d_j \qquad \ldots\ c_l\ [2]$$
$$d_{T_i} \geq d_{S_i} + 1 - M + u_i^+ M \qquad \ldots\ c_l\ [3]$$
$$d_{T_i} \geq d_{S_i} + 1 - M + u_i^- M \qquad \ldots\ c_l\ [4]$$
$$d_j \leq M \qquad \ldots\ c_l\ [5]$$

**Text S2:** Additional parameter settings and the inverse CARNIVAL pipeline

It should be noted that, while the objective function is able to rank network solutions according to the pre-defined criteria as described in the manuscript, this does not imply that the best-scoring network solutions from each parameter setting need to be similar. We therefore also explored the results generated from multiple $\alpha$-to-$\beta$ ratios. While larger node penalties frequently did not produce solutions, a node penalty between 0.03 and 1.5 resulted in similar results and similar performance for standard CARNIVAL. Given that inverse CARNIVAL was found to be more sensitive towards changes in node penalty, a value between 0.03 and 0.5 is recommended for inverse CARNIVAL. Summarised results from the study of multiple $\alpha$-to-$\beta$ ratios can be found in Supplementary Text S3.

The inverse CARNIVAL pipeline allows network inference without protein target information. While the ILP formulation itself requires known target proteins, a list of all potential perturbed nodes can be directly generated from the prior knowledge network, and can be used to capture all potential paths. In this study, we generated this list from all nodes with only outgoing and no incoming edges.

In the original ILP formulation, input nodes are not subjected to the node penalty and were already pre-defined with regard to their activity state. In inverse CARNIVAL, even if the activity state of the inputs are unknown, an input node penalty is still desired to limit the number of input nodes in the solution network. This can be achieved by adding an extra perturbation node to the prior knowledge network which has both outgoing activating and inhibiting edges to all potential input nodes. This perturbation node then serves as an unpenalized target of perturbation and can be linked to all potential input nodes without restricting their activity state. These potential input nodes are then penalized as normal additional nodes if deregulated.

**Text S3.** Summarised results from the study of multiple $\alpha$-to-$\beta$ ratios.

### 1. Standard CARNIVAL

While the original study by Melas *et al.* chose parameters in a way that one fitted measurement justifies up to five additional nodes ($\alpha/\beta = 5$), it is unclear how strongly changes in the node penalty $\beta$ affect the solutions obtained and why this particular ratio was selected. To evaluate the sensitivity of the method to changes in this penalty ratio, the mismatch penalty was set to $\alpha = 5$ and the effect of changes in node penalty tested around the parameter values of $\beta = [0; 2]$.

As expected by its definition, it was found that the number of nodes and interactions dropped with increasing node penalty (Figure ST1). In contrast, the overall interaction to node ratio and the number of models did not show a clear trend (results not shown). Only 45% of the perturbations (9 out of 20) solutions were derived from CPLEX with a node penalty of $\beta = 2$. Comparison between the solutions derived with each node penalty in each perturbation revealed that the solutions changed with modified node penalties. However, for the standard CARNIVAL benchmarking results, similar node penalties lead to similar results and the sign is not contradicting over all considered node penalties for 96.9% (3,955/4,082) of the nodes. This indicates that the node penalty has an effect which might result in alternative, but rarely contradicting solutions.

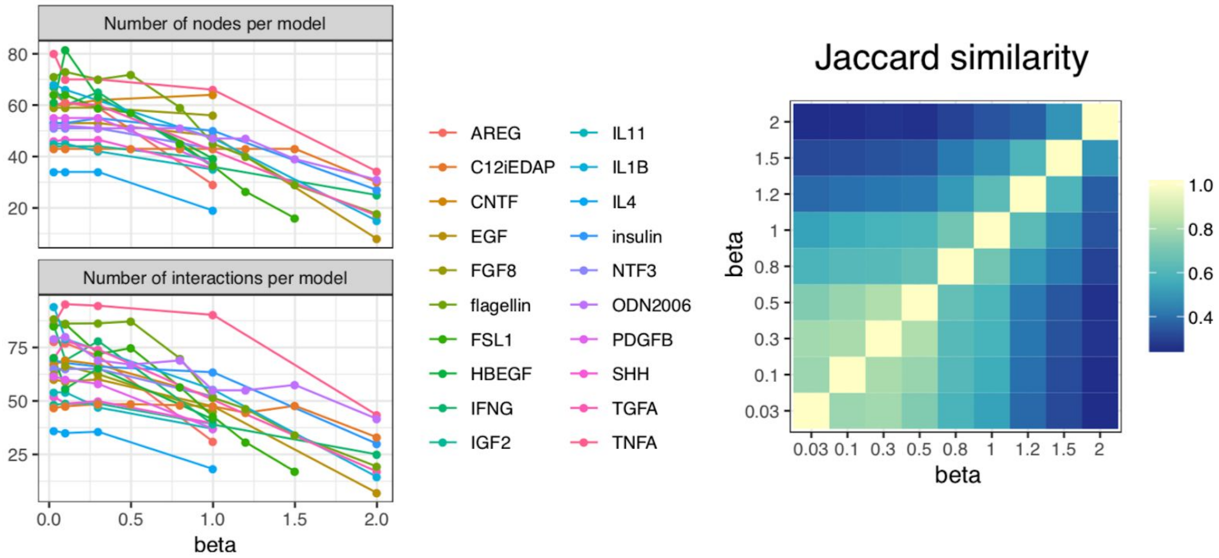


**Figure ST1: Effects of changes in node penalty β.** Overall, the number of nodes and interactions decreases with increasing node penalty, while the number of models and the interaction to node ratio do not show consistent trends. The Jaccard Index between the union of signed nodes obtained with different β is generally high between similar node penalties and decreases with increasing changes in β. Only the solutions with TF weights are compared in the figures, but similar trends were observed without TF weights.

To evaluate the effect of modifications on the node penalty β with respect to the ability of CARNIVAL to capture upstream alterations, the inferred node activities were first compared with the experimental phosphoprotein levels. However, only 3-4 of these phosphoproteins were predicted as dysregulated by CARNIVAL on average and could be matched with an activity inferred from experimentally measured phosphoprotein levels. Due to the limited availability of phosphoprotein levels, meaningful statistical testing could not be performed. As an alternative approach, the performance was evaluated based on the agreement with expected KEGG pathways with a two-step inference approach instead (see Methods).

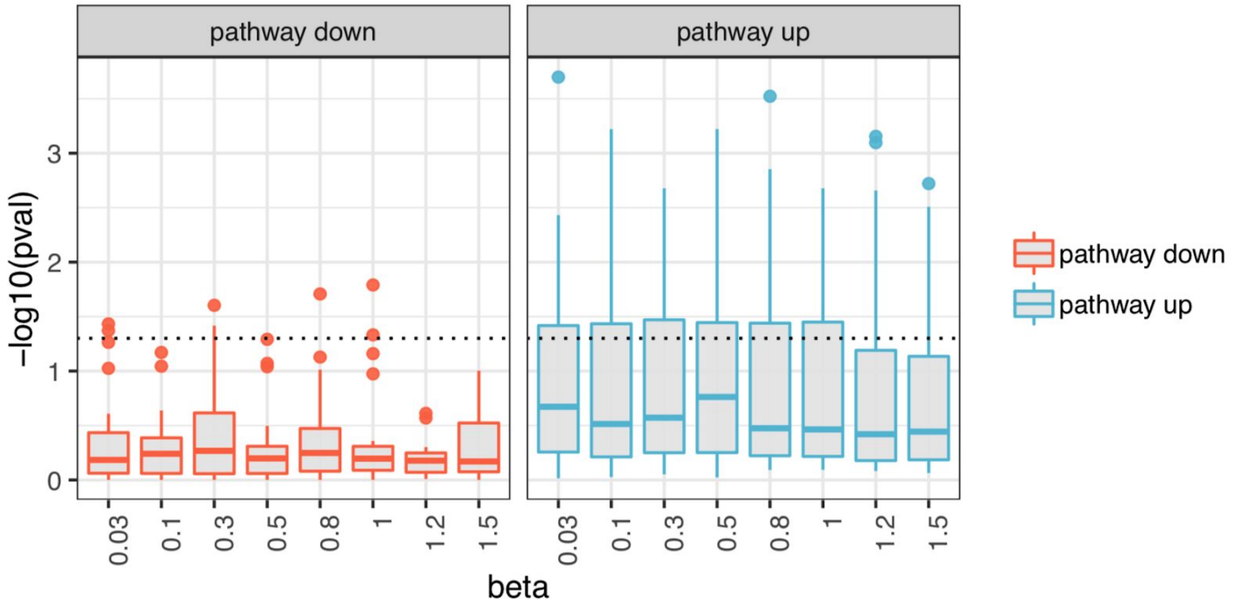


**Figure ST2: Enrichment of the perturbation-attributed pathway set in dysregulated pathways inferred from CARNIVAL over different node penalty β.** The KEGG enrichment of the attributed pathway set in the deregulated pathways shows overall similar results over different node penalty values. Generally, the attributed pathway set is slightly more enriched in activated than in inhibited pathways. The figure compares only solutions with TF weights, but similar trends were observed without TF weights.

Only minor fluctuations are observed in the enrichment of the perturbation-attributed pathway set over different node penalties and a general trend or optimum is not noticeable. Additionally, it should be noted that the perturbation-attributed pathway set is slightly more significantly enriched in activated than inhibited pathways (Figure ST2), which means that the expected perturbation-attributed pathways are more enriched in activated pathways. For CARNIVAL with known perturbation, results with a node penalty of β = 1 are selected for display given that no improvement can be observed by deviating from the original mismatch to node penalty ratio of Melas *et al.*

### 2. Inverse CARNIVAL

As a starting point, inverse CARNIVAL was first run with the previously used TF weights and a node penalty of β = 1. However, this resulted in networks with zero similarity to the previous solutions with known targets of perturbation in all cases. Further investigations revealed that all of the derived nodes are TFs inferred from DoRothEA and at the same time in the list of potential input nodes (Figure ST3). Given that these nodes only possess outgoing but not incoming reactions they can only appear as input nodes, which explains why zero similarity between inverse and standard CARNIVAL was observed.

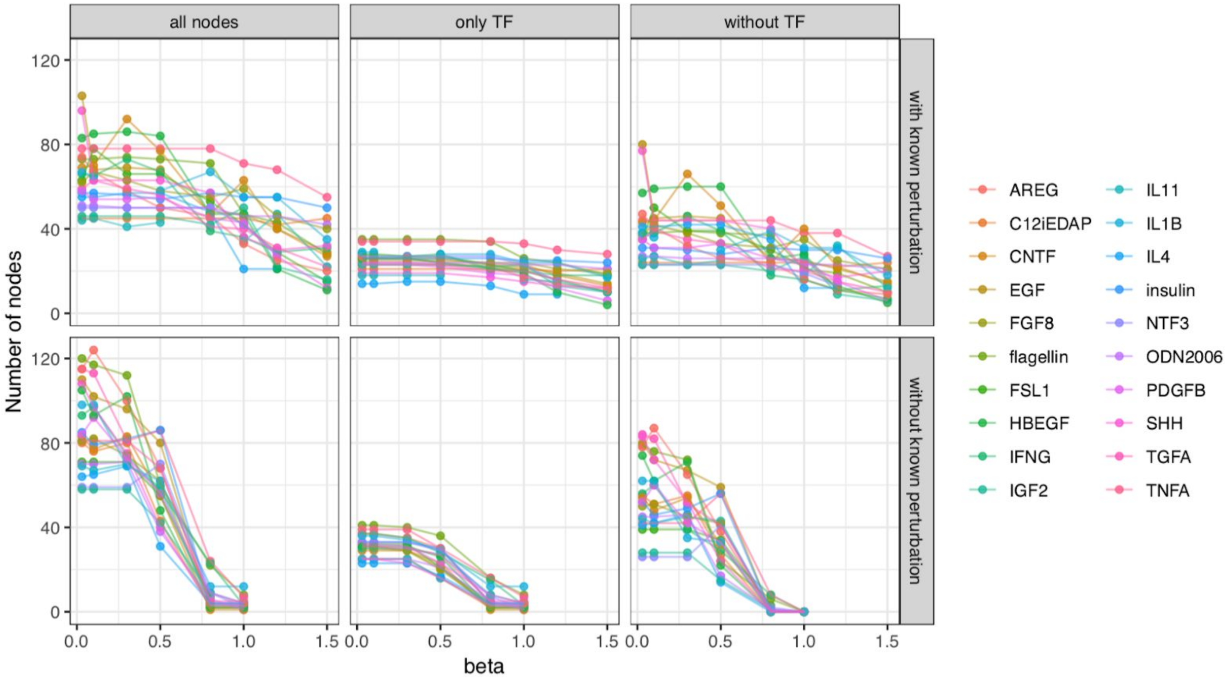


**Figure ST3: Effects of changes in node penalty β on solution nodes.** Without known perturbation, the inverse CARNIVAL pipeline is more sensitive towards changes in β. With a node penalty of 1, all appearing nodes are TFs predicted by DoRothEA, which are by definition not penalized.

In Figure ST3, the effect of the node penalty was also screened for inverse CARNIVAL and showed that the number of nodes dropped more rapidly with increasing node penalty. Also its solutions were hence more sensitive to adaptations in β than the ones of CARNIVAL with known perturbation. Given that clear trends or optima were again not identifiable by a comparison of KEGG enrichment performance over β (results not shown), a node penalty of β = 0.5 was chosen for further benchmarking given that this resulted in similar numbers of nodes compared to CARNIVAL with known perturbation.

**Text S4. Fluorescence immunohistology.**

For fluorescence immunohistology we used 3 biopsies collected from healthy renal transplantation donors and compared with 3 biopsies collected for primary diagnosis from IgAN patients. Patients had active IgAN to be included as demonstrated by IgA deposition in glomeruli. Renal needle biopsy specimens were initially collected in PBS to remove blood contamination upon biopsy. Washed tissue specimens were then fixed in normal formalin (48h) and subsequently embedded in paraffin blocks and sectioned on a microtome at a thickness of 10um. Sections were then deparaffinized and antigen-unmasked using a heat-mediated antigen retrieval buffer (Abcam) for 2h at $100^0$C. Sections were washed 3 times, blocked in 10% donkey serum in PBS (1h) and incubated with appropriate primary antibodies (mouse anti-RhoA; and rabbit anti beta-catenin; both from Abcam. Goat anti-IgA1 from Sigma). Primary antibodies were used at 1:100 dilution in 10% donkey serum and incubated with sections for 16h. Sections were washed 3 times in PBS-tween and incubated for 2h using donkey anti-mouse (AlexaFluor 647), donkey anti-rabbit (AlexaFluor 568) and donkey anti-goat (AlexaFluor 488) IgGs. Finally, sections were washed 3 times and mounted using VectaShield mounting solution (H1000). Images were taken on a Zeiss Axioscope coupled to a TissueGnostics imaging suite and a 20x lens. The same channel settings were used for all imaged sections (100ms exposure for 488 and 400ms exposure for 568 and 647).
